# Dynamic nanocomplexes of antibiotics and biofilm-degrading enzymes overcome biofilm tolerance to treat *Pseudomonas aeruginosa* infections

**DOI:** 10.64898/2026.11.16.767167

**Authors:** Alaleh Yourdkhani, Noah Y. Brittain, Chinekwu Nwagwu, Joon H. Lee, Jingwen Cao, Sara Jamshid-Nezhad, Joel A. Finbloom

## Abstract

*Pseudomonas aeruginosa* biofilms limit antibiotic penetration and protect embedded bacteria, making established infections difficult to treat. To combat the biofilm, administering tobramycin (Tob) with deoxyribonuclease I (DNase) could pair antimicrobial action with biofilm matrix degradation, but their distinct physicochemical properties hinder co-delivery and transport into the biofilm core. Taking advantage of the acidic biofilm microenvironment, we developed pH-responsive nanocomplexes (NCs) that expand, favoring local retention of the NCs and their cargo. We used stepwise electrostatic precomplexation to co-load Tob and DNase into the NCs, then electrostatically added a PEG-bearing polymer to their surface to improve NC distribution within biofilms, forming PEG-NCs. Across established biofilms of three *P. aeruginosa* strains, these dynamic NCs overcame biofilm-associated tolerance toward tobramycin and produced the most consistent reductions in bacterial burden and biomass. Formulation-function analyses coupled with biofilm imaging suggested complementary contributions from PEG and active DNase components. In an established murine abscess model, a single intra-abscess PEG-NC injection reduced bacterial burden by approximately 2 log-fold and improved abscess healing, whereas dose-matched Tob and Tob+DNase controls did not significantly improve infection outcomes. Overall, dynamic NCs mediate the co-delivery of biofilm matrix disruptors with antibiotics to improve treatment of established biofilm infections.

## 1. Introduction

Antimicrobial resistance is an escalating global health threat, with an estimated 1.14 million deaths directly attributable to bacterial resistance in 2021. These rates are projected to increase by 2050 to 1.91 million directly attributable deaths and over 8 million associated deaths annually [1]. Among resistant pathogens, *Pseudomonas aeruginosa* is particularly difficult to eradicate, as it can acquire antibiotic resistance, adapt metabolically, and form biofilms. Biofilm growth is a major mechanism of antibiotic tolerance, in which bacteria such as *P. aeruginosa* (PA) are encased in a matrix of extracellular polymeric substances (EPS) composed of polysaccharides, proteins, lipids, and extracellular DNA (eDNA) [2–4]. The EPS composition and spatial organization vary among PA strains and within individual biofilms, creating heterogeneous microenvironments that influence antimicrobial transport and bacterial metabolic state [3–6]. Recalcitrant biofilm-associated PA infections are prevalent in cystic fibrosis lung infections, as well as skin and soft tissue infections such as abscesses and chronic wounds, where repeated antibiotic exposure often fails to clear the infection [7–10].

Tobramycin (Tob) is an aminoglycoside antibiotic commonly prescribed for *P. aeruginosa* infections [10]. Although tobramycin is active against planktonic PA, its activity is markedly reduced against PA biofilms. The positive charge of tobramycin associates with negatively charged biofilm matrix components, such as eDNA, delaying its transport and producing uneven exposure across the biofilm [11,12]. In addition to acting as a physical barrier, eDNA can chelate divalent cations and contribute to local acidification, inducing bacterial responses related to aminoglycoside resistance [12–14]. The widespread presence of eDNA across PA biofilms [2–4], together with its effects on tobramycin transport and aminoglycoside resistance, makes eDNA a clinically relevant matrix target to improve tobramycin activity within established biofilms.

Deoxyribonuclease I (DNase) degrades eDNA, and recombinant human DNase is used clinically as a mucolytic in cystic fibrosis [15]. DNase-mediated matrix disruption can improve antibiotic access to biofilm-associated bacteria, but requires accompanying antibacterial activity to clear surviving or dispersed bacterial cells [16–19]. Maintaining DNase activity and access to extracellular DNA is therefore an important delivery objective [20,21], alongside co-delivery with antibiotics such as tobramycin. Since matrix sequestration can limit antibiotic penetration to deeper biofilm regions [11,12,30], coordinating matrix disruption with antibiotic delivery may help improve exposure of embedded bacteria.

Nanoparticles have shown promise for their ability to encapsulate and deliver cargo such as antibiotics [22–24]. Furthermore, engineering nanoparticle properties such as size, surface chemistry, and stimuli-responsive behavior can direct biofilm-particle interactions to improve drug distribution and antimicrobial efficacy [23,25–29]. Previous nanoparticle delivery strategies have improved tobramycin transport or combined antibiotics with DNase in biofilm *in vitro* models and cystic fibrosis patient sputum [30–32]. Spatial pH gradients in *P. aeruginosa* biofilms [38] may provide a local cue for NC expansion, potentially favoring retention of co-loaded Tob and DNase within the biofilm. However, no dynamic approaches that respond to biofilm microenvironmental cues, such as pH, have been reported for tobramycin-DNase nanoparticle co-delivery, nor has it been evaluated using in vivo infection models. We have previously leveraged polyelectrolyte nanocomplexes, self-assembling nanoparticles formed through electrostatic association of oppositely charged components, as carriers to co-deliver tobramycin and antimicrobial silver for PA infection treatment; however, our previous NC formulations did not display dynamic stimuli-responsive properties [22].

In this work, we developed pH-sensitive tobramycin-DNase co-loaded polyelectrolyte nanocomplexes (NCs) to overcome biofilm-mediated antibiotic tolerance and treat *P. aeruginosa* infections. To facilitate biofilm-responsive behavior, we formulated NCs that combined electrostatic cargo loading with pH-sensitive polyelectrolytes and a poly(ethylene glycol) (PEG) surface layer to reduce nonspecific matrix adhesion during initial biofilm penetration. We hypothesized that, upon biofilm penetration, pH-responsive NCs would swell in acidic biofilm regions, promoting physical confinement and local retention of NCs within biofilms, while co-releasing tobramycin and DNase to enhance antibiofilm activity (**Figure 1**). We evaluated both PEGylated and non-PEGylated NC formulations against established *P. aeruginosa* biofilms of the reference strains PAO1 and PA14, as well as the clinical isolate LESB58. PEGylated tobramycin-DNase nanocomplexes (PEG-NCs) overcame biofilm-associated antibiotic tolerance and produced the most consistent reductions in biofilm bacterial burden and biomass. In a murine model of an established abscess PA infection, PEG-NCs reduced bacterial burden by ∼2 log-fold and promoted abscess healing, representing a significant improvement over dose-matched non-formulated tobramycin or tobramycin-DNase combinations, which showed no measurable improvement. Taken together, this work demonstrates the development of a new pH-responsive nanocomplex drug carrier capable of dynamic co-delivery of antibiotics with enzyme adjuvants for the treatment of otherwise tolerant *P. aeruginosa* biofilm infections.

**Figure 1.**
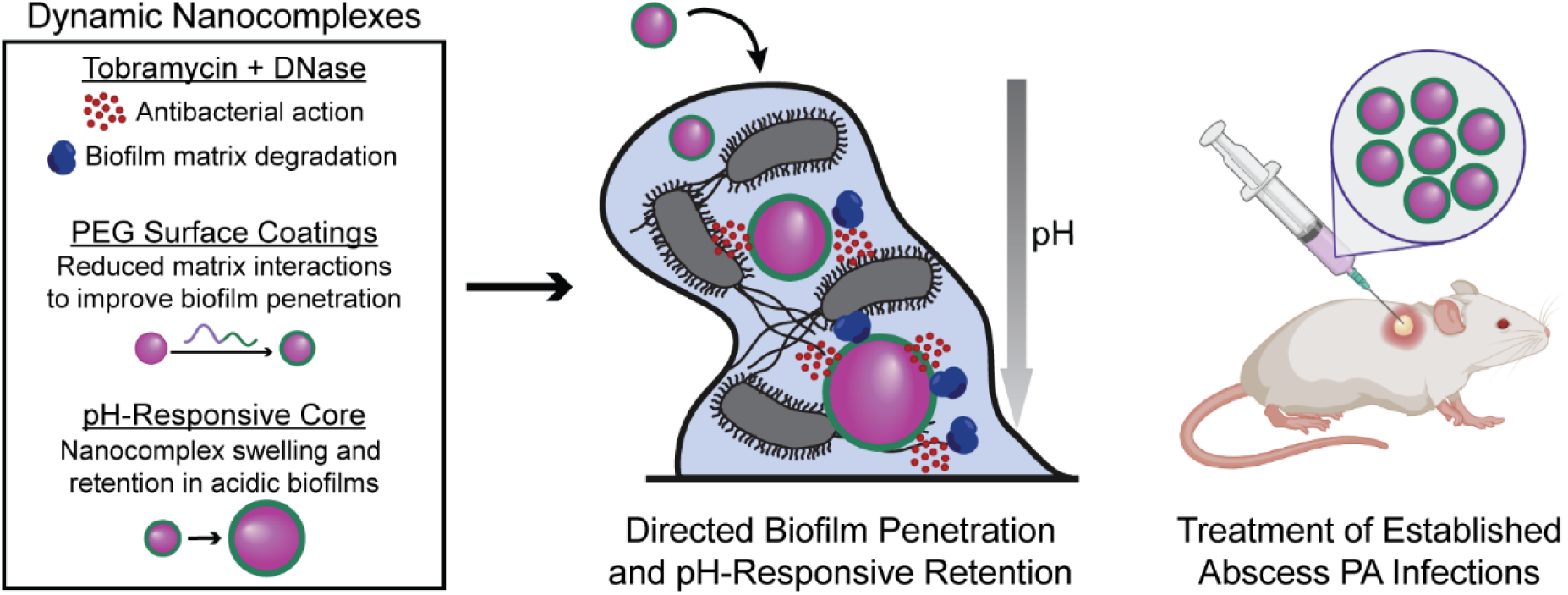
Schematic overview of dynamic nanocomplexes for tobramycin-DNase co-delivery. PEGylated, pH-responsive nanocomplexes reduce matrix interactions and improve distribution through established *P. aeruginosa* biofilms, followed by acid-triggered nanocomplex swelling, retention, and local cargo release. DNase-mediated matrix degradation and tobramycin-mediated bacterial killing together enhance biofilm treatment and improve clearance of established abscess infection in vivo.

## 2. Results and Discussion

### 2.1. Co-Loading of Tobramycin and DNase into Polyelectrolyte Nanocomplexes

Tobramycin and DNase were co-loaded into polyelectrolyte NCs through a stepwise electrostatic assembly strategy in which each cargo was first paired with an oppositely charged polymer (**Figure 2A**). The chemical structures and molecular weights of the formulation components are shown in **Table S1**. Tobramycin has reported pKa values ranging from 6.2 to 8.6, giving it an overall positive charge at the formulation pH of 7.5 [33]. We therefore screened for a pH-sensitive anionic polyelectrolyte capable of forming a uniform tobramycin precomplex with a target hydrodynamic diameter below 500 nm, selected to be below reported biofilm pore dimensions [34]. Preliminary evaluation of several anionic polyelectrolytes against these criteria led to the selection of poly(methyl vinyl ether-alt-maleic acid) (PMVEMA) (**Table S1**). PMVEMA contains two carboxylic acid groups per repeat unit, with reported pKa values of approximately 3.5 and 6.5 [35]. At pH 7.5, these groups are predominantly deprotonated, providing the negative charge required for electrostatic complexation with tobramycin. DNase required a complementary loading strategy. At pH 7.5, negatively charged DNase was paired with poly-L-lysine (PLL), building on our previous use of PLL-mediated DNase loading onto polystyrene nanoparticles [20]. The resulting DNase-PLL precomplex was combined with the anionic tobramycin-PMVEMA precomplex to form NCs containing both cargoes (**Figure 2A** and **Table S1**).

**Figure 2.**
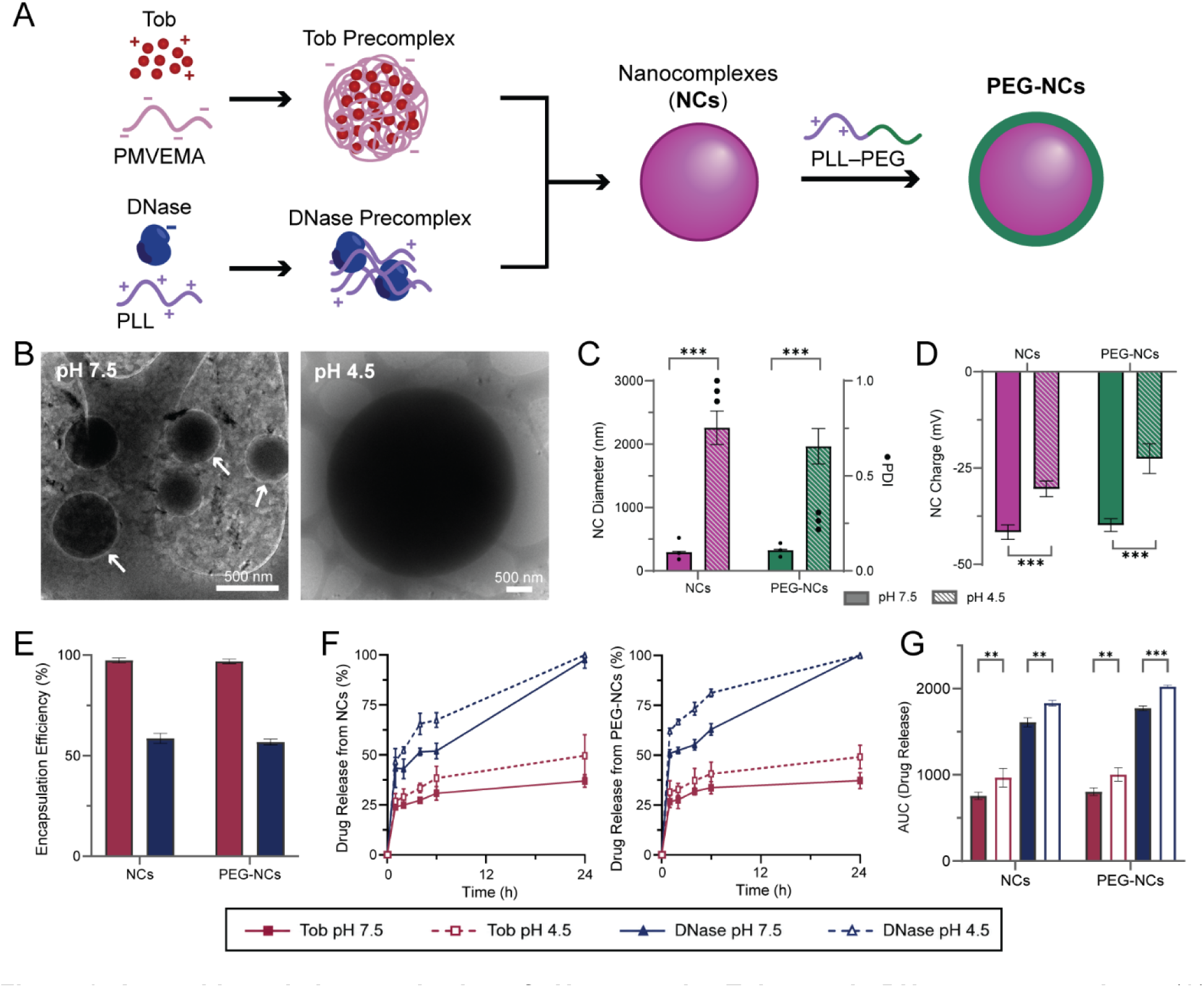
Assembly and characterization of pH-responsive Tobramycin-DNase nanocomplexes. (A) Stepwise assembly of NCs and PEG-NCs. (B) Representative cryo-TEM images of PEG-NCs at pH 7.5 and 4.5; the white arrow indicates the hollow nanocomplex morphology. Scale bars = 500 nm. (C) Hydrodynamic diameter (bars) and PDI (dots) and (D) zeta potential charge measurements of NCs and PEG-NCs under neutral and acidic conditions. (E) Tob and DNase encapsulation efficiencies. (F) Cumulative Tob and DNase release over 24 h. (G) Corresponding release AUC analysis. Data are presented as mean ± SD. **P < 0.01 and ***P < 0.001.

Component ratios, polymer molecular weights, and assembly conditions were screened using a design-of-experiments approach in Minitab based on cargo incorporation, hydrodynamic diameter, polydispersity index (PDI), and zeta potential. A low PDI was favored to obtain a narrow particle size distribution for formulation uniformity, while a negative zeta potential was selected to minimize initial electrostatic attraction between NCs and anionic biofilm matrix components such as eDNA [11,12]. Based on these criteria and initial checkerboard assays to confirm DNase-tobramycin synergy (**Figure S1**), a DNase:tobramycin ratio of 1:3 was selected and an optimized NC formulation with a DNase:PLL:tobramycin:PMVEMA mass ratio of 1:0.6:3:5.6 was determined. We postulated, based on prior reports [23], that the exposed polyelectrolyte surface of NCs could promote nonspecific matrix interactions, thereby restricting transport through the biofilm prior to pH-induced NC swelling and retention. NCs were therefore coated with a final layer of poly-L-lysine-poly(ethylene glycol) block copolymer (PLL-PEG; Table S1) to provide a bioinert PEG surface and increase initial biofilm penetration [36,37]. NCs and PEG-NCs were compared throughout subsequent in vitro evaluations.

High encapsulation efficiencies were observed for both NC and PEG-NC formulations. Tobramycin encapsulation efficiencies were 97.4 ± 2.5% for NCs and 96.9 ± 1.1% for PEG-NCs, while DNase encapsulation efficiencies were 61.7 ± 1.3% and 58.8 ± 1.5%, respectively (**Figure 2E**). The higher encapsulation efficiency of tobramycin is consistent with strong electrostatic association between its high cationic charge and polyanionic PMVEMA.

NCs and PEG-NCs displayed similar physicochemical properties at pH 7.5, with hydrodynamic diameters of 293 ± 16 nm and 326 ± 13 nm, respectively, and PDI values of 0.110 ± 0.058 and 0.110 ± 0.036 (**Figure 2C**). Cryogenic transmission electron microscopy (cryo-TEM; **Figure 2B**) and conventional transmission electron microscopy (TEM; **Figure S3**) supported the dynamic light scattering (DLS) measurements. PEG-NCs also displayed a lower-contrast halo in cryo-TEM, consistent with a hydrated PEG-rich surface layer. Both NCs and PEG-NCs had negative zeta potentials, with PEG-NCs displaying a slight but not statistically significant increase in overall charge (**Figure 2D**). In separate stability measurements, PEG-NCs maintained relatively consistent hydrodynamic diameters over 72 h in PBS at 4 °C under static conditions and in PBS and LB at 37 °C under shaking (65 rpm), indicating colloidal size stability under the tested conditions (**Figure S2D**).

### 2.2. Nanocomplex pH-Responsive Swelling and Cargo Release

Having established efficient co-loading and NC formation, we next examined whether NCs and PEG-NCs responded to acidic conditions relevant to *P. aeruginosa* biofilms. PA biofilms have spatial pH gradients, with local pH values of approximately 3.5-4.5 measured within cores of biofilm microcolonies and more neutral values reported toward the microcolony exterior [38]. We therefore used pH 4.5 as an acidic, biofilm-relevant model condition. As the pH was decreased below the PMVEMA pKa of 6.5, progressive protonation of its carboxyl groups reduced the negative charge of PMVEMA and promoted NC swelling. At 15 min, hydrodynamic diameter increased to 2259 ± 265 nm for NCs and 1966 ± 281 nm for PEG-NCs (**Figure 2C**). The zeta potentials simultaneously became significantly less negative, increasing from -41.6 ± 1.9 to -30.4 ± 2.0 mV for NCs and from -39.8 ± 1.6 to -22.6 ± 3.9 mV for PEG-NCs (**Figure 2D**).

At pH 4.5, PEG-NCs also exhibited a lower PDI than unmodified NCs (**Figure 2C**), indicating a more uniform particle population during acid-induced swelling. This behavior may reflect steric repulsion between surface-associated PEG chains, which can reduce particle-particle aggregation and improve colloidal stabilization [36]. Cryo-TEM and conventional TEM imaging supported the DLS measurements, with NCs and PEG-NCs appearing enlarged under acidic conditions while retaining an approximately spherical particulate structure (**Figures 2B and S2**), indicating that acidic conditions induced substantial NC expansion and swelling without complete loss of particulate structure.

We next determined whether this physicochemical transition altered cargo release. At pH 7.5, NCs and PEG-NCs released ∼37% of their tobramycin cargo over 24 h, while at pH 4.5, tobramycin release increased to ∼50% for both formulations (**Figure 2F**). DNase exhibited considerably faster release overall, reaching complete release from both formulations at both pH values by 24 h but with faster release by 6 h under acidic conditions (**Figure 2F**). Consistent with these kinetics, area-under-curve (AUC) analysis demonstrated significantly greater tobramycin and DNase release at pH 4.5 than at pH 7.5 from both NC and PEG-NC formulations (**Figure 2G**). Thus, the acidic condition that reduced PMVEMA charge density and promoted NC swelling also altered cargo release behavior. PLL-PEG modification did not perturb pH-responsive behaviors, allowing PEG-NCs to combine a PEGylated surface with the pH responsiveness of the underlying NC structure.

To ensure DNase activity was maintained, we used a fluorescence-based double-stranded DNA (dsDNA) degradation assay adapted from a previously reported method [39]. After 40 min incubation, free DNase and PEG-NCs each produced approximately 90% DNA-signal depletion, with no significant difference between them (**Figure S4C**). This endpoint assay supported preservation of DNase function during formulation.

To evaluate the potential long-term stability and storage of our drug carriers, we tested whether PEG-NCs could be lyophilized (Lyo-PEG-NCs) and subsequently reconstituted in aqueous media. Reconstituted Lyo-PEG-NCs produced approximately 77% DNA-signal depletion in the same assay (**Figure S4C**), indicating retained but slightly reduced DNase activity relative to fresh PEG-NCs (**Figure S4A,B**). Hydrodynamic diameter and zeta potential remained comparable to fresh PEG-NCs (**Figure S4A,B**), while PDI was also comparable (**Figure S2B**). Release profiles were also similar after reconstitution (**Figure S4D**).

### 2.3. Antibiofilm Activity and Distribution of Tobramycin-DNase Nanocomplexes in Pseudomonas aeruginosa Biofilms

NCs and PEG-NCs were evaluated for activity against established *P. aeruginosa* biofilms to assess whether combining tobramycin and DNase in a single formulation improved antibiofilm activity and whether PEG coatings further affected this response.

Biofilms were treated with Tob, Tob+DNase, NCs, or PEG-NCs at tobramycin-equivalent concentrations ranging from 7 to 225 µg/ml. Colony-forming unit (CFU) enumeration measured viable bacterial burden, while crystal violet (CV) staining measured attached biomass, including bacterial cells and retained matrix. NCs and PEG-NCs produced stronger concentration-dependent reductions in bacterial burden and biomass than the free treatment groups, with the clearest separation observed in clinical strain LESB58 biofilms (**Figure 3A,B**). Across the complete concentration range, PEG-NCs produced the lowest bacterial-burden AUC for all three strains and the lowest biomass AUC for PAO1 and LESB58 (**Figure 3A,B**). NCs and PEG-NCs also produced the strongest biomass reductions in PA14, although their biomass AUCs did not differ significantly **(Figure 3B**). Despite this strain-dependent variation, PEG-NCs produced the most consistent antibiofilm response across PAO1, PA14, and LESB58 strains. The difference between free and formulated treatments was most pronounced in LESB58. At 225 µg/ml, approximately 60% of LESB58 biomass remained after Tob and Tob+DNase treatment, compared with approximately 16% and 7% after NC and PEG-NC treatment, respectively (**Figure 3B**). This is consistent with the clinical source of LESB58, which was derived from a cystic fibrosis patient with chronic PA infections and is known as a biofilm hyperproducer, promoting a robust biofilm matrix that prevents drug penetration [40]. Therefore, increased biofilm distribution and directed drug release from NCs and PEG-NCs were required to overcome LESB58 biofilm-mediated tolerance.

**Figure 3.**
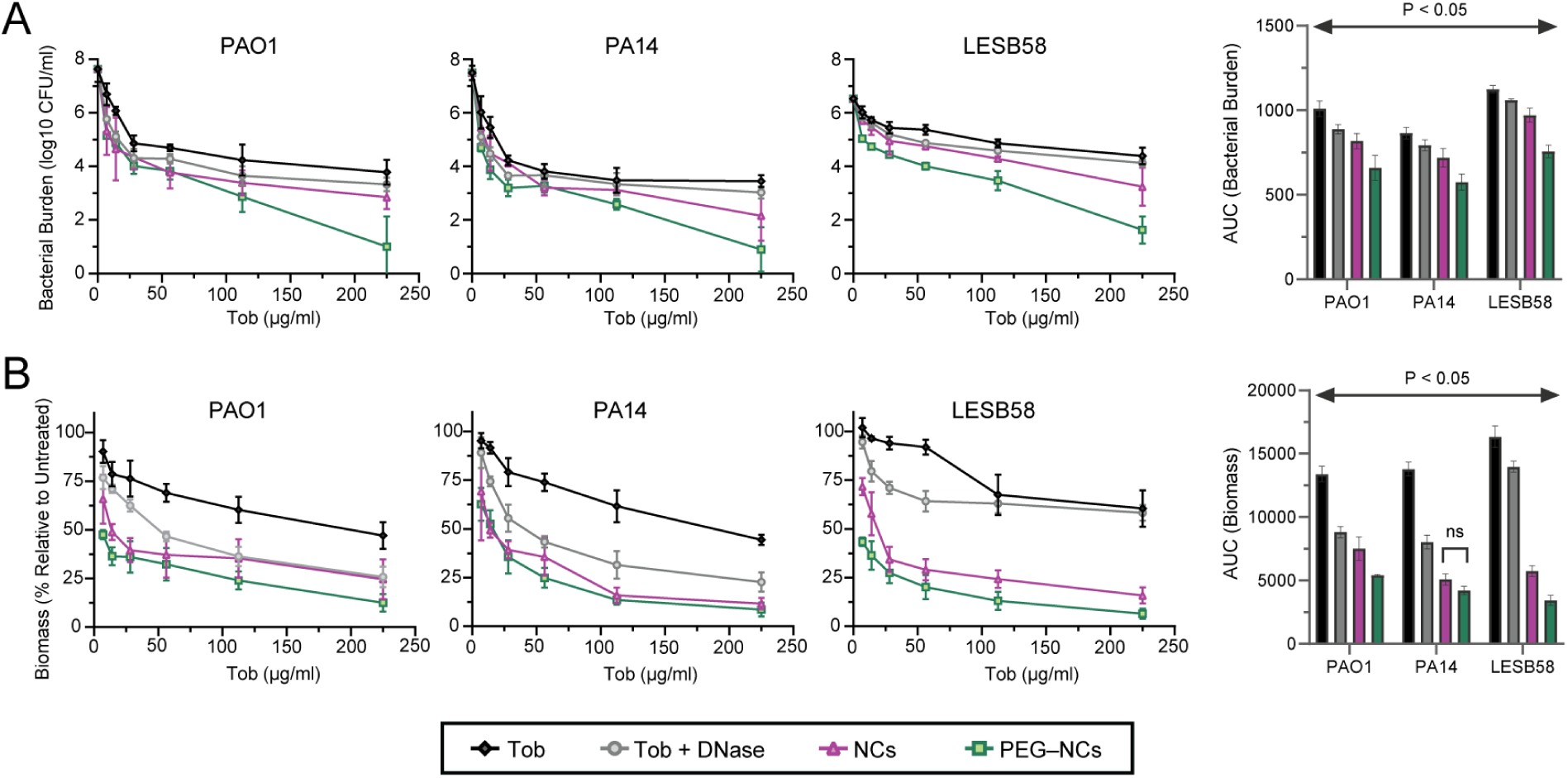
Antibiofilm activity of nanocomplexes against established *P. aeruginosa* biofilms in vitro. PAO1, PA14, and LESB58 biofilms established for 24 h were treated with increasing Tob-equivalent concentrations of Tob, Tob+DNase, NCs, or PEG-NCs. (A) Viable bacterial burden measured as log10 CFU/ml across the concentration range, with corresponding AUC analysis shown at right. (B) Biofilm biomass expressed relative to untreated biofilms across the concentration range, with corresponding AUC analysis shown at right. Data are presented as mean ± SD from independent biofilm experiments. ns, P ≥ 0.05.

We next examined whether lyophilization preserved the in vitro antibiofilm activity of PEG-NCs. Across established PAO1, PA14, and LESB58 biofilms, PEG-NCs and Lyo-PEG-NCs showed comparable concentration-dependent reductions in viable bacterial burden and biomass (Figure S4E). AUC analysis revealed no significant differences between formulations in either endpoint for PA14 or LESB58. Although Lyo-PEG-NCs produced slightly higher AUCs for PAO1 biomass (6114 ± 850 versus 5772 ± 764) and bacterial burden (686 ± 114 versus 660 ± 91), these represented relatively small changes of approximately 6% and 4%, respectively, despite reaching statistical significance. Because these modest differences were not consistently observed across strains and were not accompanied by substantial changes in NC physicochemical properties, they did not indicate a substantial or consistent loss of antibiofilm activity following lyophilization and reconstitution.

To visualize treatment-induced changes in biofilm morphology and the localization of antibacterial effects within the residual biofilm, we performed live/dead confocal microscopy. For confocal imaging, we selected a tobramycin-equivalent dose of 112.5 µg/ml (22.5 µg/ml DNase), as it produced measurable responses across all three strains while preserving sufficient residual biofilm to distinguish among treatments (**Figure 3A,B**). Representative live/dead confocal images showed reduced SYTO 9 (S9; live-associated) fluorescence and increased propidium iodide (PI; membrane-compromised/dead-associated) fluorescence following treatment with NCs and PEG-NCs (**Figure 4A**). Additionally, PEG-NCs produced the lowest S9-positive biomass across PAO1, PA14, and LESB58 (**Figure 4B**). The corresponding area-based viability index, calculated as the SYTO9-positive area divided by the PI-positive area (S9/PI), was also lowest after PEG-NC treatment, with the clearest separation observed in PAO1 and LESB58 strains (**Figure 4C**). This value is an area ratio derived from fluorescence segmentation. Therefore, it is interpreted as a relative viability index rather than a direct fraction of viable cells.

**Figure 4.**
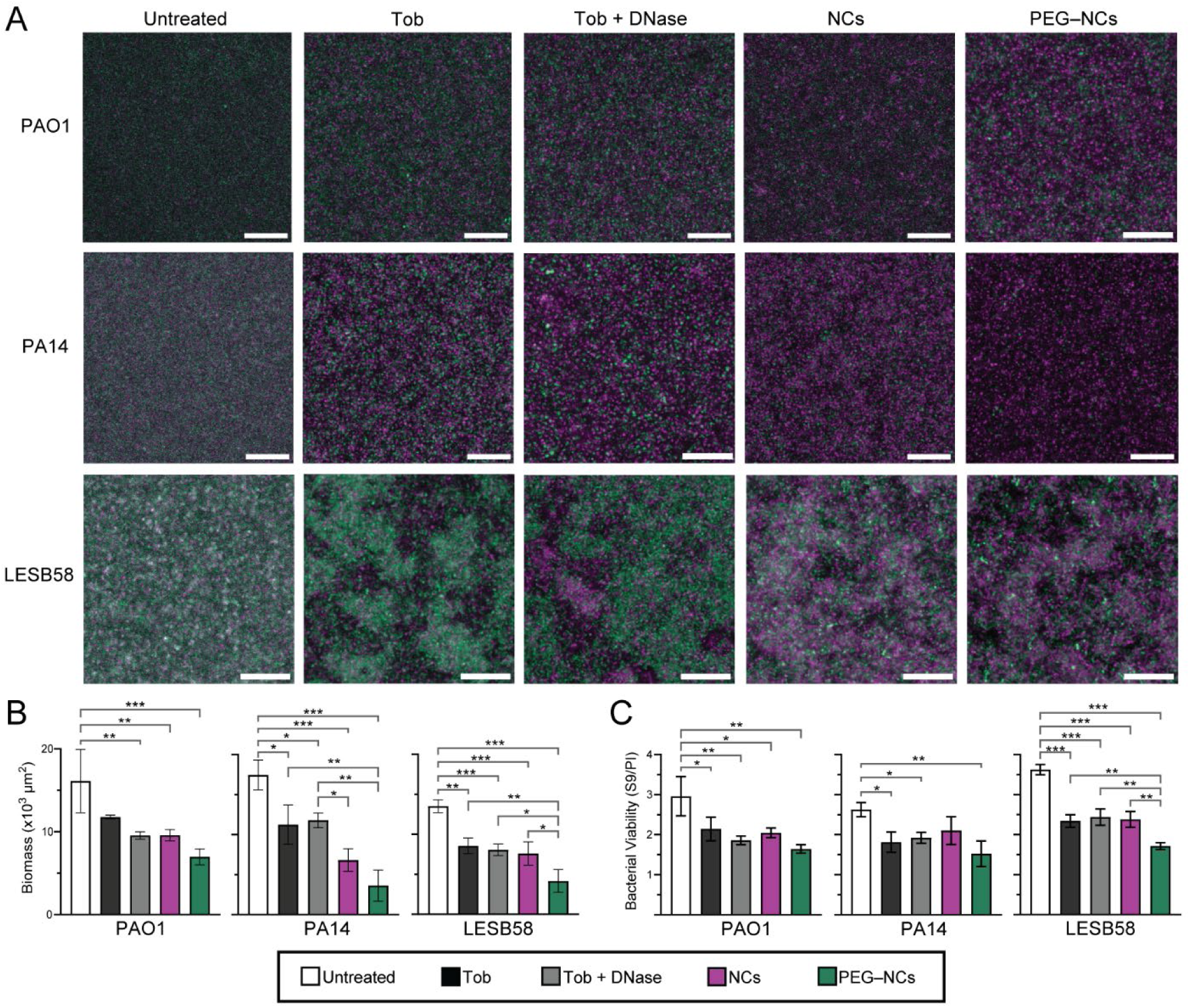
Microscopy analysis of biomass disruption and antimicrobial activity of NCs in established *P. aeruginosa* biofilms. PAO1, PA14, and LESB58 biofilms established for 24 h were treated for 24 h with Tob, Tob+DNase, NCs, or PEG-NCs. (A) Representative live/dead confocal images of live (Syto9, S9, green) and dead (propidium iodide, PI, magenta) bacteria. Scale bars = 25 μm. (B) Biofilm biomass and (C) viability index calculated as the S9-positive area divided by the PI-positive area (S9/PI). Data are presented as mean ± SD from three independent experiments. *P<0.05, **P<0.01, and ***P<0.001.

As extracellular DNA forms a core component of the *P. aeruginosa* matrix and can bind positively charged aminoglycosides such as tobramycin [12,16], we therefore compared PEG-NCs containing active DNase with formulations bearing heat-inactivated enzyme (iDNase-PEG-NCs). Heat inactivation did not produce detectable changes in NC hydrodynamic diameter or zeta potential (**Figure S5A,B**), and PDI remained comparable to PEG-NCs (**Figure S2B**). The loss of DNase activity was also confirmed in **Figure S5C,** allowing the biological comparison to focus primarily on DNase activity effects in NC formulations. We performed testingTesting in LESB58 biofilms for this and all subsequent studies, as the LESB58 clinical isolate strain displayed the most robust biofilms in our in vitro analysis. This demonstrated that, iDNase-PEG-NCs had higher bacterial-burdenbacterial burden and biomass AUCs than active PEG-NCs (**Figure S5D**). These findings support prior literature showing that DNase actively contributes to biofilm matrix degradation, thereby reducing tobramycin sequestration and increasing tobramycin-bacterial interactions for subsequent antimicrobial activity [12,13,16,27].

Having identified a functional contribution from active DNase, we next investigated whether NC PEGylation and DNase activity were also associated with greater NC distribution within established biofilms. To fluorescently track the formulations, PLL was covalently labeled with Alexa Fluor 647 prior to NC assembly, and the labeled PLL was incorporated into NC, iDNase-PEG-NC, and PEG-NC formulations. PEG-NCs showed a pronounced shift toward deeper relative z-stack positions within LESB58 biofilms after 4 h of treatment (**Figure 5A,B**). To better assess normalized distribution profiles for each formulation, distributions were divided into four relative-depth regions (0-25%, 26-50%, 51-75%, 76-100%) and compared (**Figure 5C**). Each region’s fluorescence distribution was normalized to its total signal-positive area, so that these profiles described the relative distribution of formulation-associated signals across biofilm depths, rather than absolute formulation-to-formulation comparisons. For PEG-NCs, the vast majority of biofilm distribution (89%) was observed in the deeper half of biofilms, with 36% distribution in the 51-75% depth region and 53% observed within the deepest 76-100% region. In contrast, NCs and iDNase-PEG-NCs were more evenly distributed across the biofilms, with 57% and 53% distributions in the deeper half of biofilms, respectively.

**Figure 5.**
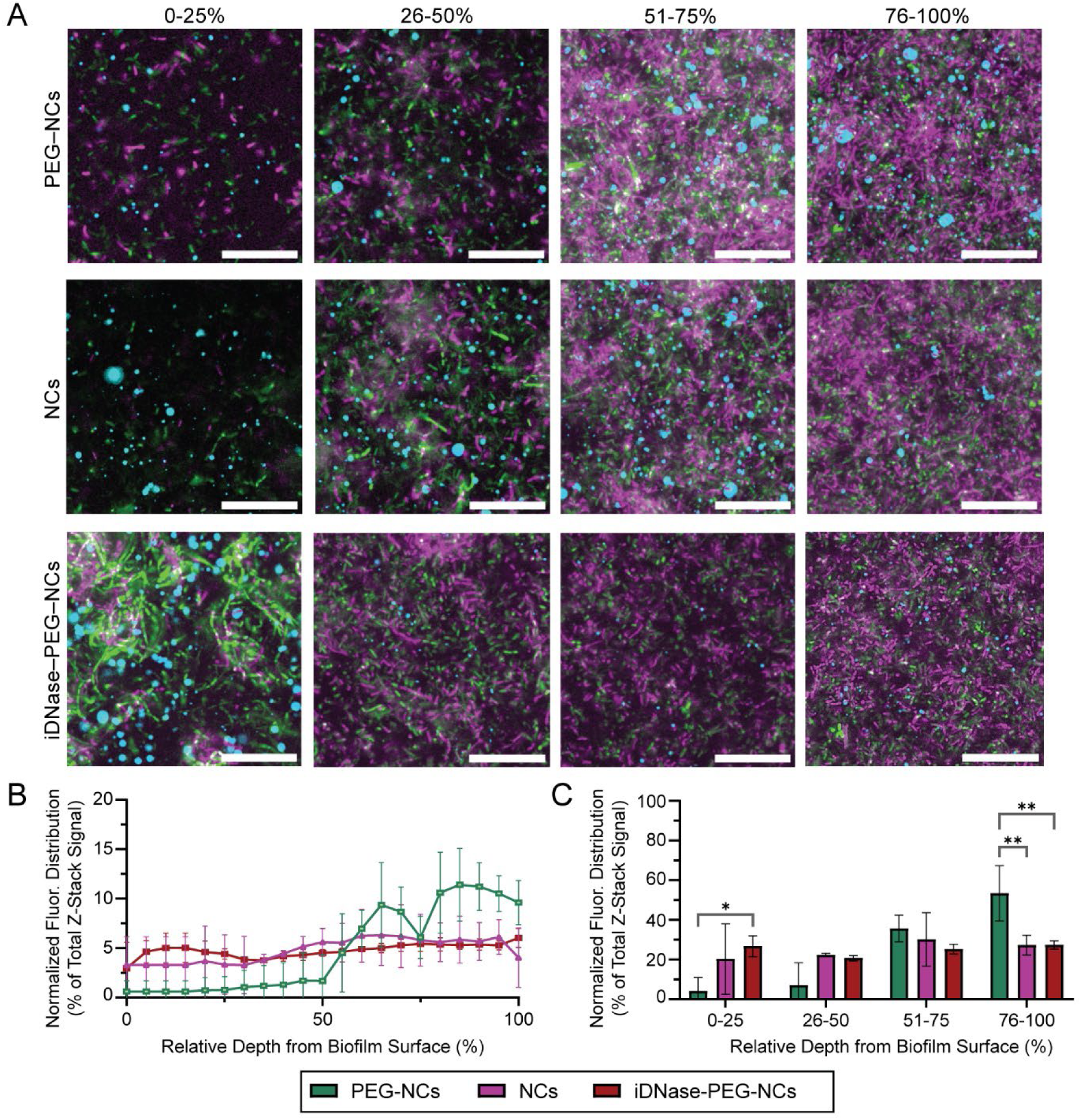
Nanocomplex distribution profiles in established LESB58 biofilms. LESB58 biofilms were grown for 24 h and then treated for 4 h with AF647-labeled PEG-NCs, NCs, or iDNase-PEG-NCs. (A) Representative 60x confocal images across four relative biofilm-z-depth regions (0–25%, 26–50%, 51– 75%, and 76–100%). AF647-labeled nanocomplexes are shown in cyan, S9-positive bacteria in green, and PI-positive bacteria in magenta, Scale bars = 25 µm. (B) Continuous relative-depth profiles of normalized AF647 occupancy. Quantitative analysis was performed using 10x confocal z-stacks, with the biofilm surface defined as 0% and the deepest acquired plane as 100%. Fluorescence distributions were normalized to the total signal within each stack, and profiles were interpolated at 5% relative-depth intervals. Data are presented as mean ± SD from three independent experiments. (C) Distribution of AF647-positive area across four relative biofilm-depth regions. *P<0.05, **P<0.01.

The different biofilm distributions between the three formulations suggests complementary functions from the formulation components. The deeper PEG-NC profile relative to NCs is consistent with prior reports that PEG can reduce adhesive or ionic interactions that would otherwise immobilize NCs within the biofilm matrix [30,37]. In a complementary fashion, iDNase-PEG-NCs retained the PEG coating without showing a comparable depth shift compared to NCs, suggesting that PEGylation alone may not fully explain the distribution of DNase-active PEG-NCs. One possibility is that PEG facilitates initial access to the biofilm, after which active DNase locally degrades biofilm matrix eDNA and permits continued redistribution of PEG-NCs through the biofilm. This hypothesis is consistent with the observed enhancement in antibiofilm activity of active PEG-NCs relative to iDNase-PEG-NCs (**Figure S5D**).

### 2.4. Cytocompatibility of NCs and PEG-NCs

We further assessed formulation cytocompatibility with mammalian cells. NIH/3T3 murine fibroblasts and HaCaT human keratinocytes were exposed to Tob, Tob+DNase, NCs, and PEG-NCs across the tested range of tobramycin-equivalent concentrations. Cell viability remained above 85% across all conditions, with broadly overlapping concentration-response profiles among the treatments (Figure S5), indicating that NC-mediated tobramycin-DNase co-delivery was well tolerated and cytocompatible.

### 2.5. Treatment of Established LESB58 Murine Abscesses with PEG-NCs

PEG-NCs were selected for in vivo evaluation as they produced the most consistent reductions in bacterial burden and biomass across the three biofilm strains. We used a localized subcutaneous PA abscess model developed by Pletzer and colleagues and subsequently applied to antimicrobial evaluation in other studies [9,41,42]. This model uses a bioluminescent LESB58 lux^+^ strain to enable noninvasive monitoring of infection localization, and generates a confined dorsal abscess with sustained bacterial burden, allowing local treatment and longitudinal monitoring [9]. Earlier studies using this model included treatment beginning 1 h after inoculation [9,41]. Here, we modified the model to treat 24 hours post-infection to better recapitulate an established biofilm infection. We delivered PEG-NCs at a concentration of 40.5 µg tobramycin and 7.5 µg DNase per mouse via a single 50 µl intra-abscess injection and the non-formulated treatments were matched to these amounts. Saline served as the vehicle control. Independent cohorts were collected at 24 and 48 h after treatment to determine treatment efficacy over time (**Figure 6A**). Bioluminescence images confirmed that LESB58 signal remained localized near the inoculation site (**Figures S7 and S8**), while bacterial burden was quantified by endpoint abscess homogenization and CFU enumeration.

**Figure 6.**
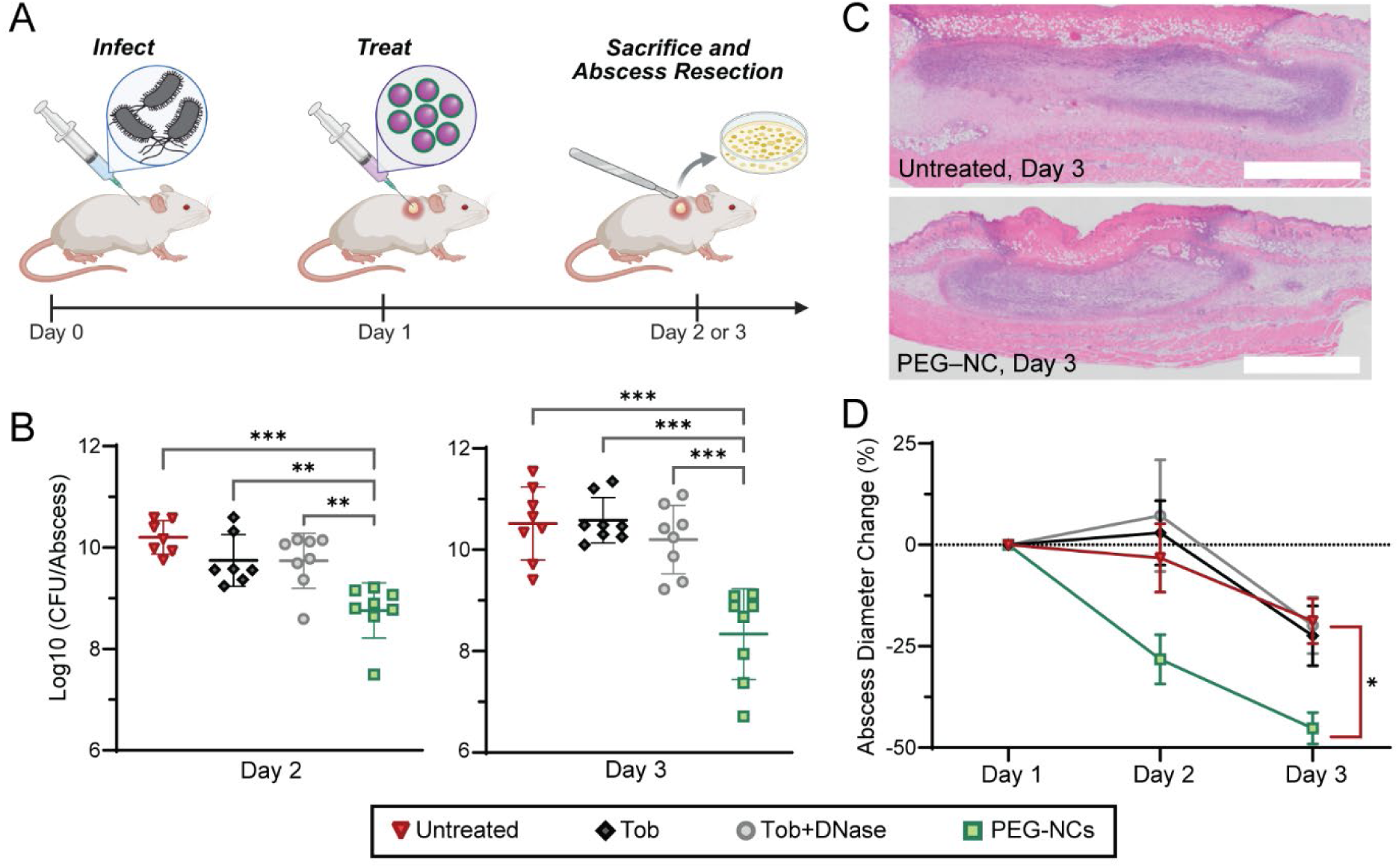
Treatment of LESB58 LUX+ murine subcutaneous abscess infections with PEG-NCs. (A) Mice were infected subcutaneously with LESB58 LUX+ on Day 0, treated with a single intra-abscess injection on Day 1, and sacrificed on Day 2 or 3. (B) Viable bacterial burden per abscess at 24 and 48 h after treatment in independent cohorts. Each point represents one mouse. (C) Representative hematoxylin and eosin (H&E)-stained sections of untreated and PEG-NC-treated abscesses collected 48 h after treatment and imaged at 10× magnification. Scale bars = 2000 μm. (D) Change in abscess diameter relative to the pretreatment measurement on Day 1 through Day 3. Data are presented as mean ± SD (n=8 mice per group for bacterial burden; n=4 per group for histology and abscess-diameter measurements). **P < 0.01 and ***P < 0.001.

At 24 h post-treatment, only PEG-NC treatment reduced bacterial burden compared to saline-treated controls, with approximately 1.3 log10 CFU/abscess reductions (**Figure 6B**). The burden was also approximately 0.9 and 1.0 log10 CFU/abscess lower than in the Tob and Tob+DNase groups, respectively, neither of which demonstrated significantly reduced bacterial burdens. A larger difference was observed in the separate cohort collected at 48 h post-treatment. PEG-NCs produced an approximately 2 log10 CFU/abscess reduction relative to saline control and were approximately 1.7-1.9 log10 CFU/abscess lower than the free treatment groups, which again did not display significant bacterial reductions relative to saline control (**Figure 6B**).

At 48 h, normalizing bacterial counts to abscess mass showed the same treatment ordering, with PEG-NCs producing the lowest CFU/mg (**Figure S8B**). Abscesses collected from PEG-NC-treated mice also had a lower average mass, although the difference was not statistically significant (**Figure S8C**). This trend may reflect reduced lesion burden, but variability in lesion size and the amount of surrounding tissue collected limited resolution of the mass endpoint. **Figure S8B-D** shows the corresponding bacterial burden, abscess mass, and body-weight measurements from the independent 24 h cohort.

Histological analysis was performed to characterize abscess lesion architecture 48 h after treatment using an independent histology cohort of mice. Representative hematoxylin and eosin (H&E)-stained sections from all groups showed a central pale or disrupted region surrounded by dense cellular tissue (**Figure 6C, S9A**), consistent with the organization reported in other murine *P. aeruginosa* abscess models [9,42]. Representative saline-, Tob-, and Tob+DNase-treated sections showed larger central lesions with extensive surrounding cellularity and disruption of adjacent tissue architecture. In contrast, the PEG-NC-treated sections appeared to contain a smaller altered region with a less extensive dense cellular area. In this cohort, we also observed abscess diameter decreases of approximately 45% by Day 3 post-PEG-NC administration, compared with approximately 20% in the other groups (**Figure 6D**). All groups initially lost body weight following infection. The PEG-NC-treated group consistently returned mouse weight closest to baseline by Day 3, whereas the other groups remained further below baseline (**Figures S7-S9**). However, the magnitude and statistical significance of the body weight recovery varied across independent cohorts, with significant differences observed in some cohorts (**Figure S7, S9**) but not others (**Figure S8**). Body weight is a systemic and relatively insensitive outcome in this localized infection model [43,44]. Small changes and inter-animal variability may therefore have limited the ability to consistently detect statistically significant differences across cohorts. Because body weight reflects overall animal condition rather than local abscess burden directly, we considered it a supportive outcome alongside other readouts.

Together with the histological analysis and reductions in bacterial burden and abscess diameter, these studies demonstrate that PEG-NC administration improved treatment outcomes in LESB58 abscess infections compared to non-formulated controls. This underscores the importance in drug carrier formulation design to direct co-delivery within biofilm infection sites.

## 3. Conclusions

We developed dynamic pH-responsive PEG-NCs that co-deliver tobramycin and DNase to treat established *P. aeruginosa* biofilm infections. Across PA strains, PEG-NCs produced the most consistent antibiofilm activity, with each component playing a key role: NC PEGylation was associated with deeper intrabiofilm distribution, active DNase contributed to matrix disruption, and acidic conditions induced NC swelling for biofilm retention and increased cargo release. In an established LESB58 PA abscess model, a single local PEG-NC dose reduced bacterial burden by approximately 2 log-fold at 48 h post-treatment, whereas dose-matched Tob and Tob+DNase did not significantly reduce burden relative to saline. Together, these findings support coordinated biofilm distribution, matrix disruption, and antibiotic-adjuvant co-delivery as a strategy for treating established *P. aeruginosa* biofilm-associated infections.

## 4. Experimental Section

### 4.1. Chemicals and Reagents

Tobramycin, PLL hydrobromide (30-70 kDa), PMVEMA (number-average molecular weight (Mn) ≈ 80 kDa; weight-average molecular weight (Mw) ≈ 216 kDa), o-phthaladehyde (OPA), fluorescein isothiocyanate isomer I (FITC), crystal violet, PLL-PEG (PEG Mn ≈ 2 kDa, PLL Mn ≈ 10 kDa), DNase I from bovine pancreas (specific activity >2000 U/mg), and Pseudomonas-selective agar were purchased from Sigma-Aldrich (St. Louis, USA). Resazurin sodium salt, Miller LB agar, cell-culture reagents, Alexa Fluor 647 NHS ester (AF647), and the LIVE/DEAD BacLight Bacterial Viability Kit containing S9 and PI were obtained from Thermo Fisher Scientific (Waltham, USA). Miller LB broth was obtained from VWR International (Radnor, USA). Chemical structures and information of NC formulation components are provided in **Table S1**.

### 4.2. Instruments

Hydrodynamic diameter and PDI were measured by DLS, and zeta potential was measured by electrophoretic light scattering using a Zetasizer Nano ZS (Malvern Instruments Ltd., Malvern, UK). Absorbance measurements were performed using a SpectraMax Plus 384 microplate reader (Molecular Devices, San Jose, USA), whereas fluorescence measurements were performed using a Synergy Mx microplate reader (BioTek Instruments, Winooski, USA). Confocal imaging was performed using a Zeiss LSM 900 Airyscan 2 confocal microscope (Carl Zeiss Microscopy GmbH, Jena, Germany). TEM imaging was performed using an LVEM5 transmission electron microscope (Delong Instruments, Brno, Czech Republic). Cryo-TEM imaging was performed using an FEI Tecnai G20 transmission electron microscope (FEI Company, Hillsboro, USA) operated at 200 kV, with vitrification performed using a Thermo Scientific Vitrobot (Thermo Fisher Scientific, Waltham, USA).

### 4.3. Cell Lines and Bacterial Strains

NIH/3T3 murine fibroblasts were obtained from the laboratory of Dr. Jayachandran N. Kizhakkedathu at the University of British Columbia (Vancouver, Canada). HaCaT human keratinocytes were purchased from Cedarlane Laboratories (Burlington, Canada). *Pseudomonas aeruginosa* PAO1 and PA14 were provided by the laboratory of Dr. Cara Haney at the University of British Columbia (Vancouver, Canada). LESB58 and bioluminescent LESB58 LUX+ were provided by the laboratory of Dr. Robert E. W. Hancock at the University of British Columbia (Vancouver, Canada) [9,41,45]. Bacterial strains were maintained as glycerol stocks at -80 °C.

### 4.4. Mice

Female CD-1 mice aged 5-6 weeks were obtained from Charles River Laboratories (USA). Animals were maintained under a 12 h light/dark cycle with ad libitum access to food and water and were provided with environmental enrichment. All animal procedures were approved by the University of British Columbia Animal Care Committee under protocol A24-0153 and were conducted in accordance with the guidelines of the Canadian Council on Animal Care.

### 4.5. Preparation of NCs and PEG-NCs

All aqueous solutions were prepared using ultrapure water unless otherwise stated. Tobramycin, PMVEMA, DNase, PLL, and PLL-PEG stock solutions were prepared at 10 mg/mL. NCs were prepared in laboratory-prepared 1x phosphate-buffered saline (PBS) by separately forming tobramycin-PMVEMA and DNase-PLL precomplexes, followed by their combination at a DNase:PLL:tobramycin:PMVEMA mass ratio of 1:0.6:3:5.6. This ratio was selected following optimization of the component ratios based on particle size, PDI, zeta potential, and component encapsulation efficiency.

Tobramycin stock solution (45 µl; 450 µg) was added to 200 µl PBS and bath-sonicated for 20 s in an ice-cooled bath sonicator. PMVEMA stock solution (84 µl; 840 µg) was added during sonication, followed by another 30 s of sonication and incubation at room temperature for 15 min. In a separate tube, PLL stock solution (9 µl; 90 µg) and DNase stock solution (15 µl; 150 µg) were sequentially added to 147 µl PBS, gently mixed by pipetting, and incubated at room temperature for 15 min. The DNase-PLL precomplex was then added to the tobramycin-PMVEMA precomplex during 5 s of bath sonication in an ice-cooled sonicator, followed by incubation at room temperature for 15 min. PEG-NCs were prepared by incorporating 75 µg PLL-PEG, corresponding to a DNase:PLL-PEG mass ratio of 1:0.5. The PBS volume was adjusted accordingly to maintain a final formulation volume of 500 µl.

Following nanocomplexation, NCs and PEG-NCs were dialyzed against 1x PBS for 16 h at 4 °C using Float-A-Lyzer devices with a 300 kDa molecular-weight cutoff (MWCO; Repligen, Waltham, USA) to remove unbound components. All reported physicochemical characterization and loading measurements were performed after dialysis, and formulations were used immediately after dialysis unless otherwise stated.

Lyo-PEG-NCs were prepared by flash-freezing and lyophilizing PEG-NCs, followed by reconstitution to the original formulation volume using Milli-Q water. iDNase-PEG-NCs were prepared by heating DNase at 75 °C for 20 min; all subsequent preparation and dialysis steps were identical to those used for PEG-NCs. Tobramycin-PMVEMA control precomplexes were prepared using 450 µg tobramycin and 840 µg PMVEMA as described above, with the final volume adjusted to 500 µl using PBS.

### 4.6. Physicochemical Characterization

Hydrodynamic diameter and PDI were measured by DLS, and zeta potential was measured by electrophoretic light scattering. Hydrodynamic diameter was reported as the Z-average diameter calculated by cumulant analysis of the intensity autocorrelation function. Measurements were performed at 25 °C after 2 min of equilibration. Three independently prepared formulation batches were analyzed, with three measurements collected and averaged for each batch.

The pH-responsive behavior of the formulations was evaluated by mixing 100 µl of each formulation with 100 µl of either 1x PBS adjusted to pH 7.5 or 200 mM sodium acetate adjusted to pH 4.5. Samples were incubated for 15 min at 37 °C before measurement. Hydrodynamic diameter and PDI were measured directly following incubation. For zeta-potential measurements, samples were diluted 1:100 in 2 mM KCl immediately before analysis.

Colloidal stability of PEG-NCs was evaluated over 72 h PBS at 4 °C (static) and in PBS or LB at 37 °C with shaking at 65 rpm. Hydrodynamic diameter was measured at 25 °C at the indicated time points.

Particle morphology was examined by TEM and cryo-TEM. For TEM, 5 µl of sample was applied to glow-discharged TEM grids and imaged at 5.5 kV. For cryo-TEM, 3 µl of sample was applied to glow-discharged 300-mesh lacey F/C grids and vitrified with a blot time of 1 s and a blot force of 1. Vitrified grids were transferred under cryogenic conditions and imaged at 200 kV without staining or additional sample processing.

### 4.7. Cargo Encapsulation

Tobramycin and DNase encapsulation efficiencies were determined by quantifying the unencapsulated cargo remaining in the supernatant after formulation and dialysis. NCs, PEG-NCs, Lyo-PEG-NCs, and iDNase-PEG-NCs were centrifuged at 16,900 x g for 10 min at 24 °C. Encapsulation efficiency (%) was calculated as [(initial cargo - unencapsulated cargo)/initial cargo] x 100%.

Tobramycin was quantified using an o-phthaladehyde (OPA) fluorescence assay as previously described [22,46]. OPA reagent was prepared at 1 mg/mL in 100 mM sodium borate buffer (pH 10.4) containing 10% v/v methanol and 0.5% v/v β-mercaptoethanol. OPA reagent, isopropanol, and sample or tobramycin standard were combined at a 1:6:1 volume ratio and incubated for 20 min at room temperature. Fluorescence was measured at 360/450 nm (excitation/emission, ex/em) using matrix-matched calibration curves.

DNase was quantified using FITC-DNase as a fluorescent tracer [20]. DNase (10 mg/ml) was reacted with FITC at a 15:1 FITC:DNase molar ratio in 0.1 M sodium bicarbonate buffer (pH 8) for 3 h. For 1 ml of DNase solution, 45 µl of 90 mM FITC in dimethyl sulfoxide (DMSO) was added. Unconjugated FITC was removed by dialysis against water for 6 h at 4 °C using a 3 kDa MWCO membrane (Thermo Scientific, Waltham, USA). The FITC-to-DNase labeling ratio was determined using the Proteins & Labels application on a NanoDrop spectrophotometer (Thermo Fisher Scientific, Wilmington, DE, USA) from DNase absorbance at 280 nm and FITC absorbance at 494 nm, with correction for the contribution of FITC to protein absorbance, yielding an estimated ratio of 2.4 FITC molecules per DNase molecule. FITC-DNase was detected at 488/520 nm (ex/em) using matrix-matched calibration curves.

### 4.8. Cargo Release Measurements

For all release studies, 1x PBS at pH 7.5 or 20 mM sodium acetate containing 137 mM NaCl at pH 4.5 was used as the release medium.

For tobramycin release, 100 µl of each formulation was loaded into a 7 kDa MWCO Slide-A-Lyzer dialysis cassette (Thermo Scientific, Waltham, USA), immersed in 14 ml release medium, and incubated at 37 °C with shaking at 65 rpm. At each time point, 70 µl of the external medium was collected and replaced with an equal volume of fresh medium. Tobramycin concentrations were measured using the OPA assay described above.

For DNase release, NCs and PEG-NCs containing FITC-DNase were mixed 1:1 v/v with release medium to a final volume of 6 ml and incubated at 37 °C with shaking at 65 rpm. At each time point, 280 µl was collected and centrifuged at 16,900 x g for 20 min at 4 °C. FITC-DNase in the supernatant was quantified using pH-matched calibration curves, and the collected volume was replaced with fresh medium.

Release studies were performed in three independent experiments. In each experiment, one formulation batch was tested in three separate release vessels, and the three values were averaged.

### 4.9. DNase Enzymatic Activity

DNase activity was evaluated using the QuantiFluor dsDNA System (Promega, Madison, USA), adapted from a fluorescence-based DNase assay described previously [39]. Free DNase, NCs, and PEG-NCs were normalized to 0.2 U/µl, corresponding to 0.1 mg/mL DNase. Each sample had a total volume of 25 µl and contained 1 U DNase and 0.5 µg of the dsDNA supplied with the kit. Samples were incubated for 40 min at 37 °C. Intact dsDNA was quantified according to the manufacturer’s instructions using matrix-matched calibration curves. DNA degradation was calculated relative to a DNA-only control and expressed as a percentage of the initial DNA content. Lyo-PEG-NCs and iDNase-PEG-NCs were evaluated using the same procedure.

### 4.10. Biofilm Formation and Treatment

Static biofilms were established using a modified 96-well microtiter plate assay [47]. Bacterial cultures were inoculated directly from glycerol stocks into LB and grown overnight at 37 °C with shaking at 185 rpm. Overnight cultures were diluted 1:8 in fresh LB and grown for another 1-3 h to an optical density at 600 nm (OD600) of 0.4-0.6. Cultures were then diluted in LB to an OD600 of 0.01, transferred to 96-well plates at 100 µl per well, and incubated without agitation at 37 °C for 24 h in a humidified incubator.

Following biofilm formation, the entire 100 µl of planktonic culture was carefully aspirated from each well using a multichannel pipette without disturbing the attached biofilm. Each well was gently washed with 100 µl sterile PBS, then aspirated. Biofilms were treated with 100 µl per well of Tob, Tob+DNase, NCs, PEG-NCs, or assay-specific control formulations at the concentrations indicated for each experiment. Formulations were normalized to their tobramycin- and DNase-equivalent concentrations, and all treatment solutions were prepared in LB. Because the formulation stocks were prepared in PBS, vehicle-control wells received LB containing the same volume of PBS as the corresponding formulation-treated wells. Where indicated, Lyo-PEG-NCs were reconstituted to their original volume and evaluated alongside freshly prepared PEG-NCs.

Following treatment, plates were incubated without agitation at 37 °C for another 24 h. Three independently established and treated biofilm experiments were analyzed. Technical-replicate wells within each independent experiment were averaged before analysis.

### 4.11. Biofilm-Associated Bacterial Viability

The treatment medium was aspirated, and each biofilm was washed once with 100 µl sterile PBS. Biofilms were collected from each well using a sterile swab. The swab was transferred to 1 ml LB and sonicated for 4 min to resuspend the recovered bacteria. Suspensions were serially diluted 1:10 in sterile PBS to a final dilution of 10^-8^, and 5 µl droplets from each dilution were spot-plated in triplicate onto Pseudomonas-selective agar., and each dilution was spot-plated in triplicate onto LB agar using 5 µl per spot. Plates were incubated at 37 °C for 24 h before colonies were counted. Viable bacterial burden was expressed as log10 CFU/ml.

### 4.12. Biofilm Biomass

The entire 100 µl of treatment medium was aspirated from each well using a multichannel pipette, and each biofilm was gently washed with 100 µl sterile PBS. Plates were allowed to dry, then stained with 0.1% w/v crystal violet in ultrapure water for 20 min. Excess stain was removed, and wells were washed twice with PBS. The retained crystal violet was solubilized in 70% (v/v) ethanol, and absorbance was measured at 595 nm. Background absorbance from bacteria-free wells containing the corresponding treatments after the wash steps was subtracted, and biofilm biomass was expressed relative to untreated biofilms.

### 4.13. Confocal Live/Dead Imaging and Image Analysis

Biofilms were prepared as described above and treated with Tob, Tob+DNase, NCs, or PEG-NCs at tobramycin- and DNase-equivalent concentrations of 112.5 and 22.5 µg/mL, respectively. Untreated biofilms and single-dye controls were included. The staining stock was prepared by mixing 190 µl PBS, 5 µl PI (20 mM stock), and 5 µL S9 (5 mM stock). Each well received 96 µl treatment solution and 4 µl staining stock, giving a final volume of 100 µl and final concentrations of approximately 20 µM PI and 5 µM S9. Samples were incubated at 37 °C for 24 h while protected from light.

Images were acquired using a 20x objective on a ZEISS LSM 900 Airyscan 2 microscope with ZEN 3.4 software. S9 and PI were excited using 488 and 561 nm lasers, respectively. Acquisition settings were kept constant across treatment groups within each strain. Each z-stack contained 21 optical sections acquired at 2.5 µm intervals over a total depth of 50 µm. In each independent experiment, one field of view was acquired from each of three technical-replicate wells per treatment group. Three independent biofilm experiments were performed. Representative panels shown in Figure 3 were center-cropped to 50% of the original 20x field of view for presentation without resizing, interpolation, or sharpening. All quantitative analyses were performed on the original uncropped 20x z-stacks.

Confocal z-stacks were analyzed using a custom Python pipeline. Each z-plane was converted to floating-point intensity and smoothed with a Gaussian filter (σ = 1.0 pixel) before segmentation, while unblurred intensities were retained for intensity measurements. S9-positive biomass was segmented independently in each plane using Otsu thresholding after clipping intensities at the 99.5th percentile to limit the influence of hot pixels. PI-positive area was segmented using a fixed rule applied identically to all samples: the robust PI Otsu threshold was multiplied by 1.8, and for sparse planes in which S9-positive biomass covered <15% of the field, the threshold was constrained to be at least the 99th percentile of PI intensity outside the S9 mask plus 1 intensity unit. No per-slice adaptive scale search or biomass-guided threshold lowering was used. Biofilm biomass was quantified as S9-positive area, and the viability index shown in Figure 3 was calculated as A_S9/A_PI, where A_S9 and A_PI are the thresholded S9- and PI-positive areas over the full field of view. The reciprocal A_PI/A_S9 was retained as the area-based death-fraction metric. Plane-level measurements were averaged across each z-stack, and the three technical-replicate wells were then averaged to obtain one value for each independent biofilm experiment. Analyses were performed in Python 3 using NumPy, scikit-image, pandas, matplotlib, and aicsimageio/aicspylibczi for .czi Zeiss inputs.

### 4.14. Nanocomplex Biofilm Distribution and Relative-Depth Analysis

NCs, PEG-NCs, and iDNase-PEG-NCs were evaluated for their distribution within established LESB58 biofilms. For fluorescence imaging, PLL was labeled with Alexa Fluor 647 (AF647) N-hydroxylsuccinimide (NHS) ester before preparing the formulations. PLL (10 mg/ml) was reacted with AF647 NHS ester at a 1 mol% dye-to-amine ratio in 0.1 M sodium bicarbonate buffer (pH 8.3). For 1 ml of PLL solution, 0.478 µmol AF647 NHS ester was dissolved in anhydrous DMSO and added dropwise to PLL during sonication. The reaction was rotated overnight at room temperature while protected from light. Absolute AF647 was labeling was not independently quantified; instead, formulations were normalized to equal concentrations and comparable input AF647 fluorescence was confirmed across formulations before treatment.

AF647-PLL was then used to prepare NCs, PEG-NCs, and iDNase-PEG-NCs for NC-biofilm distribution experiments. Formulations were normalized to equal concentrations, and comparable input AF647 fluorescence was confirmed across formulations before treatment. Established LESB58 biofilms were treated with the labeled formulations for 4 h as well as S9 and PI, and imaged using consistent acquisition settings within each series. Quantitative relative-depth analysis was performed using 10x confocal z-stacks, whereas representative higher-magnification optical sections were acquired using a 60x objective. Optical sections were acquired at 2.5 µm z-intervals.

AF647-associated distribution was quantified using a custom Python image-analysis pipeline. Each z-plane was smoothed using a Gaussian filter (σ = 1.0 pixel), and AF647-positive regions were initially identified using Triangle thresholding. Binary masks were opened using a 1-pixel-radius disk, and connected components smaller than 16 pixels were removed. To limit classification of diffuse background as formulation-associated signal, planes in which the preliminary mask occupied >5% of the field of view or had a mean mask intensity <6-fold above the unmasked background were re-thresholded using Yen’s method and, when required, a robust median-plus-MAD background threshold. A stack-level threshold floor was also applied to prevent isolated low-threshold planes from producing artificial signal peaks.

For each optical section, AF647 occupancy was calculated as the AF647-positive area divided by the total field-of-view area. For relative-depth analysis, the designated biofilm exterior surface within each individual z-stack was assigned a depth of 0% and the deepest acquired plane a depth of 100%. Each optical section was then assigned a relative depth based on its z-position within its corresponding stack. Individual profiles were interpolated at 5% relative-depth intervals to enable comparison and averaging across stacks with different imaging depths. Each interpolated profile was normalized so that the sum of its values equaled 100%, allowing comparison of the relative distribution of formulation-associated signal across biofilm depth rather than absolute formulation uptake.

For the regional analysis, the normalized profiles were divided into four relative-depth regions: 0-25%, 26-50%, 51-75%, and 76-100%. Values within each region were summed to determine the proportion of the total normalized stack distribution associated with that depth range. Three selected 10x image stacks were analyzed per formulation.

### 4.15. Mammalian Cell Culture and Biocompatibility

NIH/3T3 murine fibroblasts and HaCaT human keratinocytes were cultured in high-glucose Dulbecco’s modified Eagle medium (DMEM) supplemented with 10% fetal bovine serum (FBS) and 1% penicillin-streptomycin. Cells were maintained at 37 °C in a humidified atmosphere containing 5% CO2. For biocompatibility experiments, cells were seeded into 96-well plates at 10,000 cells per well in 100 µl complete medium and allowed to adhere for 24 h.

Tob, Tob+DNase, NCs, and PEG-NCs were diluted in complete medium to tobramycin-equivalent concentrations ranging from 56.25 to 450 µg/mL using twofold serial dilutions. DNase-containing treatments were adjusted to the corresponding DNase-equivalent concentrations. The culture medium was replaced with 100 µl of treatment-containing medium per well.

Following 20 h of treatment, 100 µl of resazurin solution (10% v/v in water) was added to each well and incubated for another 5 h at 37 °C while protected from light. Fluorescence was measured at 560/590 nm (ex/em). Cell-free wells containing the corresponding treatments and resazurin were included for background correction. Cell viability was expressed relative to untreated cells. Three independent experiments were performed, with three technical-replicate wells included for each condition. Technical replicates from each experiment were averaged before analysis.

### 4.16. Murine Subcutaneous Abscess Model of Biofilm-Associated Infection

The murine subcutaneous abscess model was adapted from a previously described method [9]. LESB58 lux^+^ was cultured overnight in LB at 37 °C with shaking at 185 rpm. The overnight culture was diluted in fresh LB and grown to an OD600 of 0.4-0.6. Bacteria were pelleted by centrifugation at 5000 rcf for 5 min and resuspended in saline at 1 x 10^9^ CFU/ml based on an OD600-CFU calibration curve. The final inoculum concentration was confirmed by CFU enumeration.

One day before infection, dorsal fur was removed by shaving followed by application of a commercial depilatory cream (Nair Cream Sensitive Formula; Church & Dwight Co., Inc., Ewing, USA) according to the manufacturer’s instructions. On Day 0, mice were anesthetized with isoflurane and inoculated subcutaneously in the right dorsal region with 5 x 10^7^ CFU of LESB58 LUX+ in 50 µl. The infection was allowed to establish for 24 h before treatment.

On Day 1, mice were randomly assigned to receive a single 50 µl intra-abscess injection of Tob, Tob+DNase, PEG-NCs, or 0.9% saline. Each mouse received a fixed amount of 40.5 µg tobramycin and, where applicable, 7.5 µg DNase. For a 25 g mouse, these fixed amounts corresponded to nominal doses of 1.62 mg/kg tobramycin and 0.30 mg/kg DNase. Tob and Tob+DNase administrations were amount-matched to the corresponding cargo contents of PEG-NCs. Separate cohorts were collected at 24 and 48 h after treatment, with eight mice per treatment group at each time point. An independent cohort of four mice per treatment group was used for histological analysis and abscess measurements at 48 h after treatment.

### 4.17. Clinical Monitoring and Abscess Diameter Measurement

Body weight and clinical condition were assessed on Day 0, immediately before treatment on Day 1, and at 24 h and 48 h after treatment on Days 2 and 3, respectively. Body-weight change was expressed as a percent relative to the Day 0 measurement. Abscess diameter was measured in the independent histology cohort using a digital caliper, and abscess-diameter change was expressed as a percent relative to the Day 1 pretreatment measurement.

### 4.18. In Vivo Bioluminescence Imaging

Bacterial bioluminescence was assessed immediately before treatment on Day 1 and at 24 and 48 h after treatment on Days 2 and 3. Mice were anesthetized with isoflurane, positioned dorsal side up, and imaged using a Lago X in vivo imaging system (IVIS, Spectral Instruments Imaging, Tucson, USA). Images were acquired and processed using Aura software with the same acquisition settings across treatment groups at each time point. Images were displayed using a linear color scale ranging from 1 x 10^5^ to x 10^5^ photons/s/cm^2^/sr, with the threshold set to 1 x 10^5^ photons/s/cm^2^/sr. IVIS images were used to visualize infection localization over time; viable bacterial burden was quantified independently by endpoint CFU enumeration.

### 4.19. Abscess Collection and Bacterial Enumeration

Following the final assessment at the corresponding 24 h or 48 h endpoint, mice were euthanized, and abscesses with surrounding tissue were aseptically excised. Excess surrounding tissue was trimmed where necessary, and the mass of each abscess was measured. Sterile PBS was added at a tissue-to-PBS ratio of 1:5 w/v (5 µl PBS per mg tissue). Samples were homogenized using a Bullet Blender Tissue Homogenizer (Next Advance, Troy, USA).

Homogenates were serially diluted 1:10 in sterile PBS to a final dilution of 10^-8^, and 5 µl droplets from each dilution were spot-plated in triplicate onto Pseudomonas-selective agar. Plates were incubated at 37 °C for 24 h before colonies were counted. Bacterial burden was expressed as CFU/abscess or CFU/mg and log10-transformed where indicated. Samples with no detectable colonies were recorded as 0 CFU.

### 4.20. Histological Analysis

The independent histology cohort was collected 48 h after treatment. Abscesses with surrounding tissue were excised and fixed in 10% neutral-buffered formalin for 24 h before transfer to 70% ethanol. Fixed tissues were submitted to a histology service for processing, including dehydration through a graded ethanol series, xylene clearing, paraffin embedding, sectioning at 5 µm, and hematoxylin and eosin (H&E) staining. Sections were imaged using a Zeiss Axioscan at 10x magnification to assess tissue architecture, abscess organization, and inflammatory-cell infiltration.

### 4.21. Statistical Analysis

Data are presented as mean ± SD unless otherwise stated. Each independently prepared formulation batch, independently established biofilm experiment, independent cell-culture experiment, or individual mouse was considered one independent replicate. Technical measurements from the same independent experiment were averaged before statistical analysis. Unless otherwise noted, all in vitro experiments were performed with three independent replicates. Sample sizes for the in vivo experiments are provided in the corresponding Methods sections.

Comparisons involving more than two groups were performed using one-way analysis of variance (ANOVA) followed by Tukey’s multiple-comparisons test. Two-group comparisons were performed using two-tailed *t*-tests. For concentration-response and release experiments, one AUC value was calculated separately for each independent replicate from the complete curve. Concentration-response AUC values were analyzed separately for each bacterial strain using one-way ANOVA followed by Tukey’s multiple-comparisons test. No statistical comparisons were performed between experiments of different strains. Longitudinal and endpoint mouse cohorts were analyzed independently and were not pooled.

Statistical analyses were performed using GraphPad Prism version 11.0.2 (GraphPad Software, Boston, USA). Statistical significance was defined as *p* < 0.05. Significance symbols followed the GraphPad Prism New England Journal of Medicine (NEJM) convention: \**p* for 0.01 ≤ *p* < 0.05, \*\**p* for 0.001 ≤ *p* < 0.01, and \*\*\**p* for *p* < 0.001; ns indicates *p* ≥ 0.05.

## 5. Acknowledgments

This work was supported by the University of British Columbia Animal Care Services Diagnostic & Research Histology Laboratory. The authors acknowledge the UBC Bioimaging Facility (RRID: SCR_021304). This research was supported by a peer-reviewed Cystic Fibrosis Canada 2025 Early Career Investigator Award to Joel Finbloom. J.A.F. acknowledges additional funding support from the Michael Smith Health Research BC Scholarship Program. A.Y. is supported by the NSERC CREATE-CITF program. N.Y.B. is supported by the NSERC Canadian Doctoral Scholarship. C.N. is supported by the CIHR Canadian Doctoral Scholarship.

## 6. Supporting Information

Supporting figures and tables S1-S9. Chemical structures and molecular characteristics of NC formulation components; Checkerboard analysis of tobramycin and DNase interactions in established biofilms; Physicochemical properties and colloidal stability of nanocomplex formulations; pH-Responsive morphological expansion of NCs and PEG-NCs; Effects of lyophilization on PEG-NC properties and antibiofilm efficacy; Effects of DNase inactivation on nanocomplex properties and antibiofilm activity; Cytocompatibility of NC formulations in HaCaT and NIH/3T3 cells; Bacterial burden and treatment outcomes in the independent 24-h murine abscess cohort; Bacterial burden and treatment outcomes in the independent 48-h murine abscess cohort; Longitudinal outcomes and histology from the independent 48-h murine abscess cohort.

