## Supporting Information for "Dynamic nanocomplexes of antibiotics and biofilm-degrading enzymes overcome biofilm tolerance to treat *Pseudomonas aeruginosa* infections"

### TABLE OF CONTENTS

**Table S1.** Chemical structures and molecular characteristics of NC formulation components ........................ S1

**Figure S1.** Checkerboard analysis of tobramycin and DNase interactions in established biofilms .................. S2

**Figure S2.** Physicochemical properties and colloidal stability of nanocomplex formulations ........................... S3

**Figure S3.** pH-responsive morphological expansion of NCs and PEG-NCs ................................................... S4

**Figure S4.** Effects of lyophilization on PEG-NC properties and antibiofilm efficacy ....................................... S5

**Figure S5.** Effects of DNase inactivation on nanocomplex properties and antibiofilm activity ......................... S6

**Figure S6.** Cytocompatibility of NC formulations in HaCaT and NIH/3T3 cells ........................................... S7

**Figure S7.** Bacterial burden and treatment outcomes in the independent 24-h murine abscess cohort .......... S8

**Figure S8.** Bacterial burden and treatment outcomes in the independent 48-h murine abscess cohort .......... S9

**Figure S9.** Longitudinal outcomes and histology from the independent 48-h murine abscess cohort ........... S10

**Supplementary References** ......................................................................................................... S11

| NC Component | Molecular Weight and Composition | Chemical Structure |
|---|---|---|
| tobramycin (Tob) | MW = 467.5 g/mol | 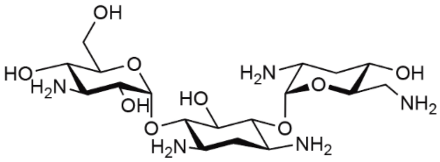 |
| deoxyribonuclease I (DNase) | MW ~ 37 kDa<br>Source: bovine pancreas<br>Specific activity > 2,000 U/mg | |
| poly(methyl vinyl ether-alt-maleic acid) (PMVEMA) | Mn ~ 80 kDa<br>Mw ~ 216 kDa | 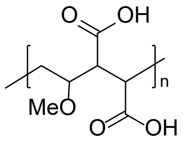 |
| poly-L-lysine (PLL) | 30-70 kDa | 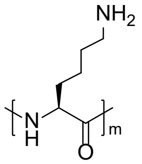 |
| methoxy-poly (ethylene glycol)-amine-poly-L-lysine (PLL-PEG) | PEG Mn 1.5-2.5 kDa<br>PLL Mn 3-8 kDa | 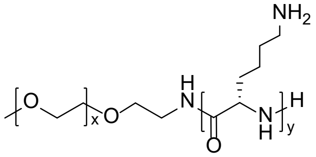 |

**Table S1. Chemical structures and molecular characteristics of components used for NC formulation.**

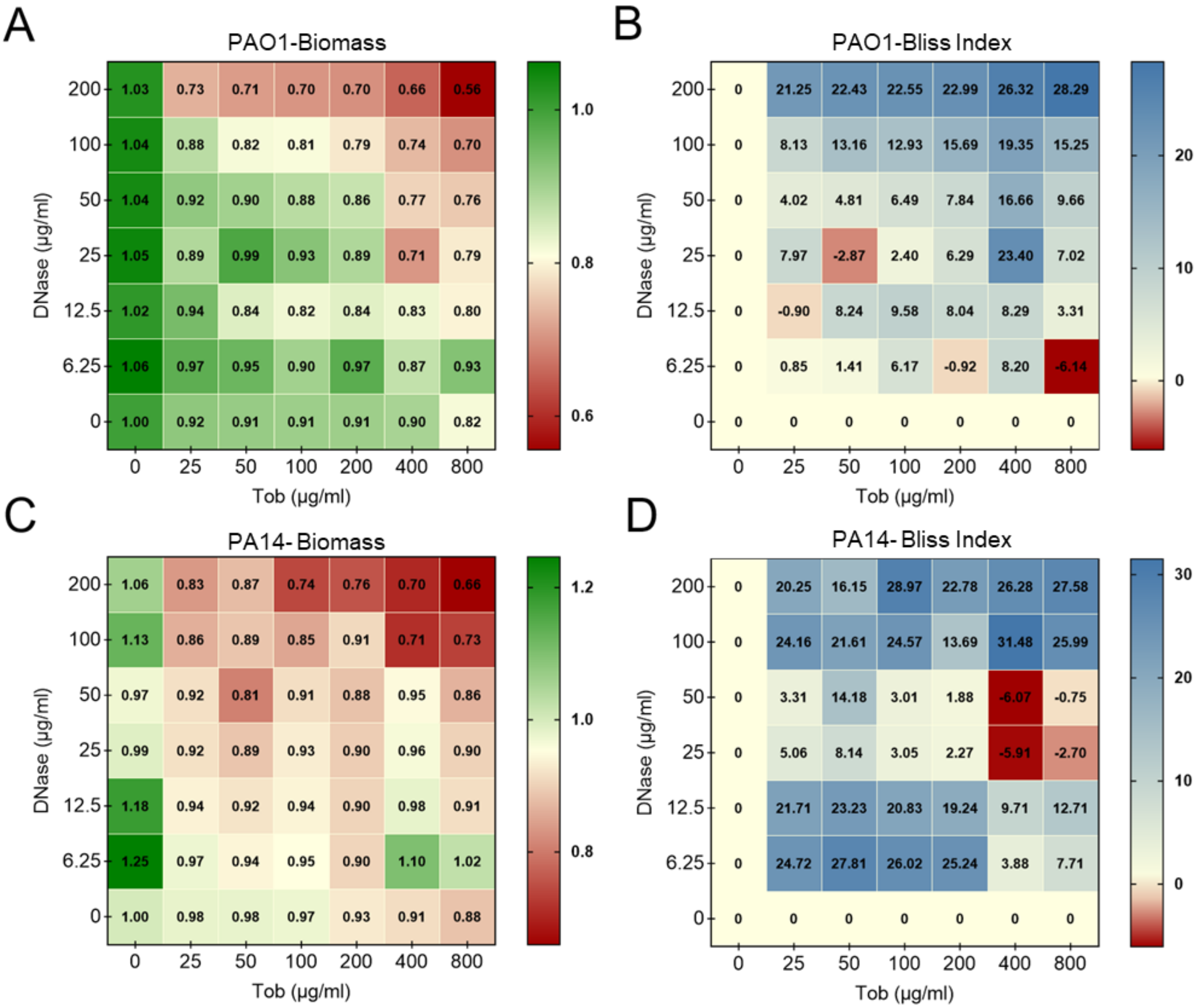


**Figure S1. Checkerboard analysis of tobramycin and DNase interactions in established *P. aeruginosa* biofilms.** PAO1 and PA14 biofilms were treated with increasing concentrations of tobramycin (Tob; 0-800 µg/ml) and DNase (0-200 µg/ml). (A, C) Biofilm biomass measured by crystal violet (CV) staining and expressed relative to untreated controls in (A) PAO1 and (C) PA14. (B, D) Corresponding Bliss excess inhibition calculated from the expected independent response and observed combination response for (B) PAO1 and (D) PA14, expressed in percentage points, following the Bliss independence model [See References S1–S3]. Positive Bliss values indicate greater biomass reduction than predicted by independent action, whereas negative values indicate less reduction than predicted. Bliss values greater than 10 are highly indicative of synergy. Data are presented as normalized mean responses from three independent experiments.

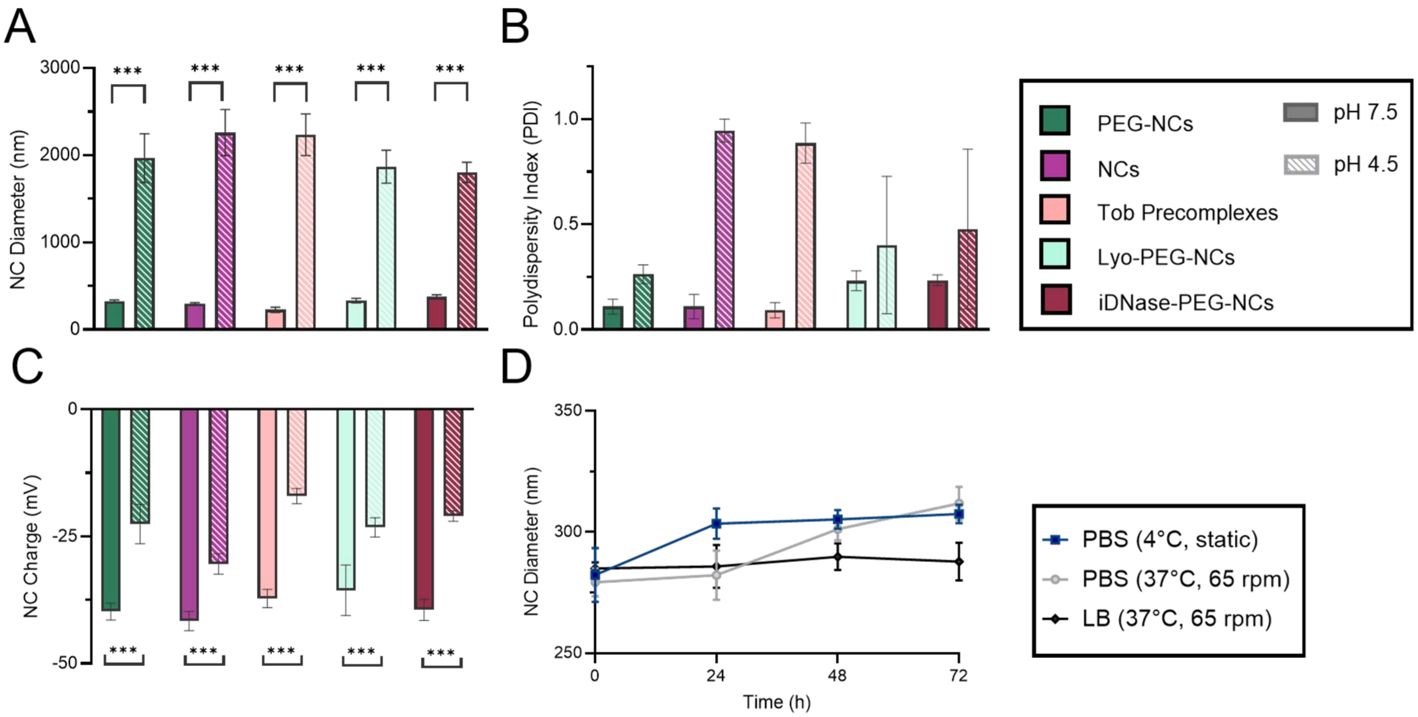


**Figure S2. Physicochemical properties and colloidal stability of nanocomplex formulations.** PEG-NCs, NCs, Tob pre-complex, Lyo-PEG-NCs, and iDNase-PEG-NCs were characterized at pH 7.5 and 4.5. (A) Hydrodynamic diameter, (B) PDI, (C) zeta potential, and (D) PEG-NCs hydrodynamic diameter over 72 h. Data are presented as mean ± SD from three independently prepared formulation batches. ***P<0.001.

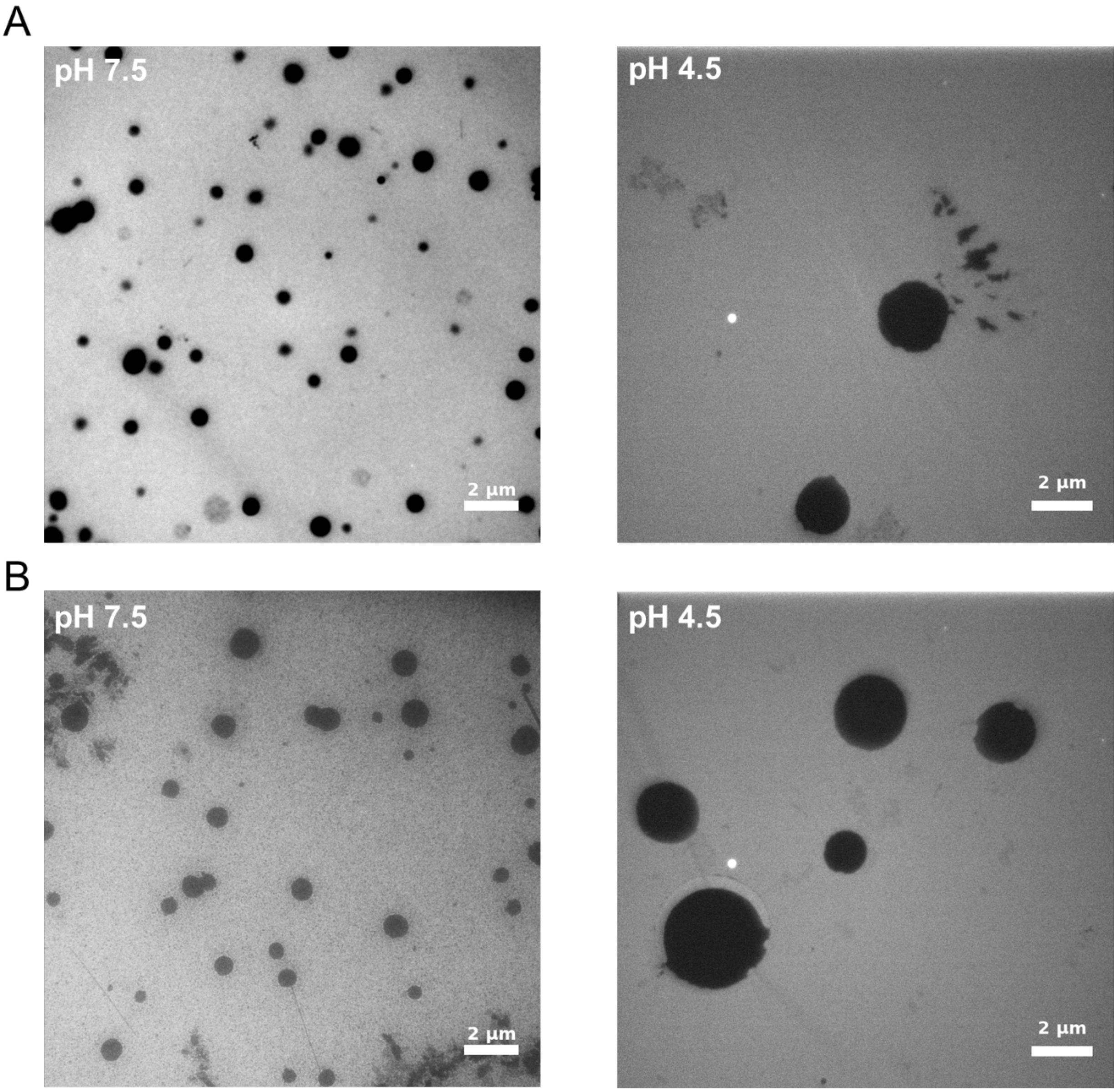


**Figure S3. PEG-NCs and NCs demonstrate pH-responsive morphological expansion.** (A) PEG-NCs and (B) NCs were imaged by conventional TEM (non-cryo-TEM) after incubation at pH 7.5 and 4.5. Representative images show increased particle size under acidic conditions. Scale bars = 2 µm.

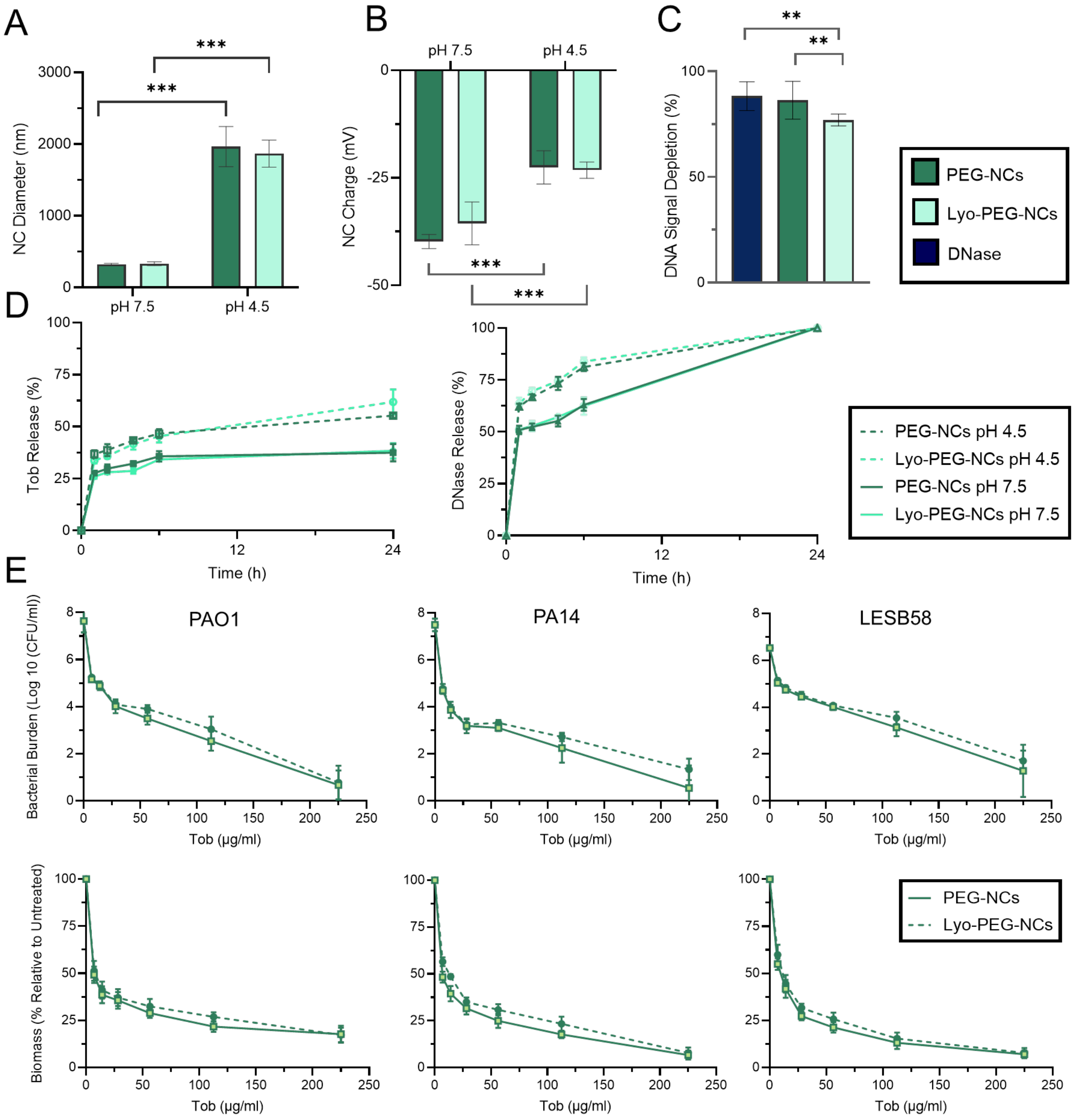


**Figure S4. Lyophilization preserves PEG-NC physicochemical properties, cargo release, DNase activity, and antibiofilm activity.** Fresh PEG-NCs and lyophilized PEG-NCs (Lyo-PEG-NCs) were compared following reconstitution. (A) Hydrodynamic diameter and (B) zeta potential at pH 7.5 and 4.5. (C) DNase activity was measured as DNA signal depletion after 40 min incubation for free DNase, PEG-NCs, and Lyo-PEG-NCs. (D) Cumulative tobramycin (Tob; left) and DNase (right) release from PEG-NCs and Lyo-PEG-NCs at pH 7.5 and 4.5 over 24 h. (E) Concentration-response profiles for viable bacterial burden (top) and biofilm biomass (bottom) in established PAO1, PA14, and LESB58 biofilms treated with PEG-NCs or Lyo-PEG-NCs. Data are presented as mean ± SD from three independent experiments. **P < 0.01 and ***P < 0.001.

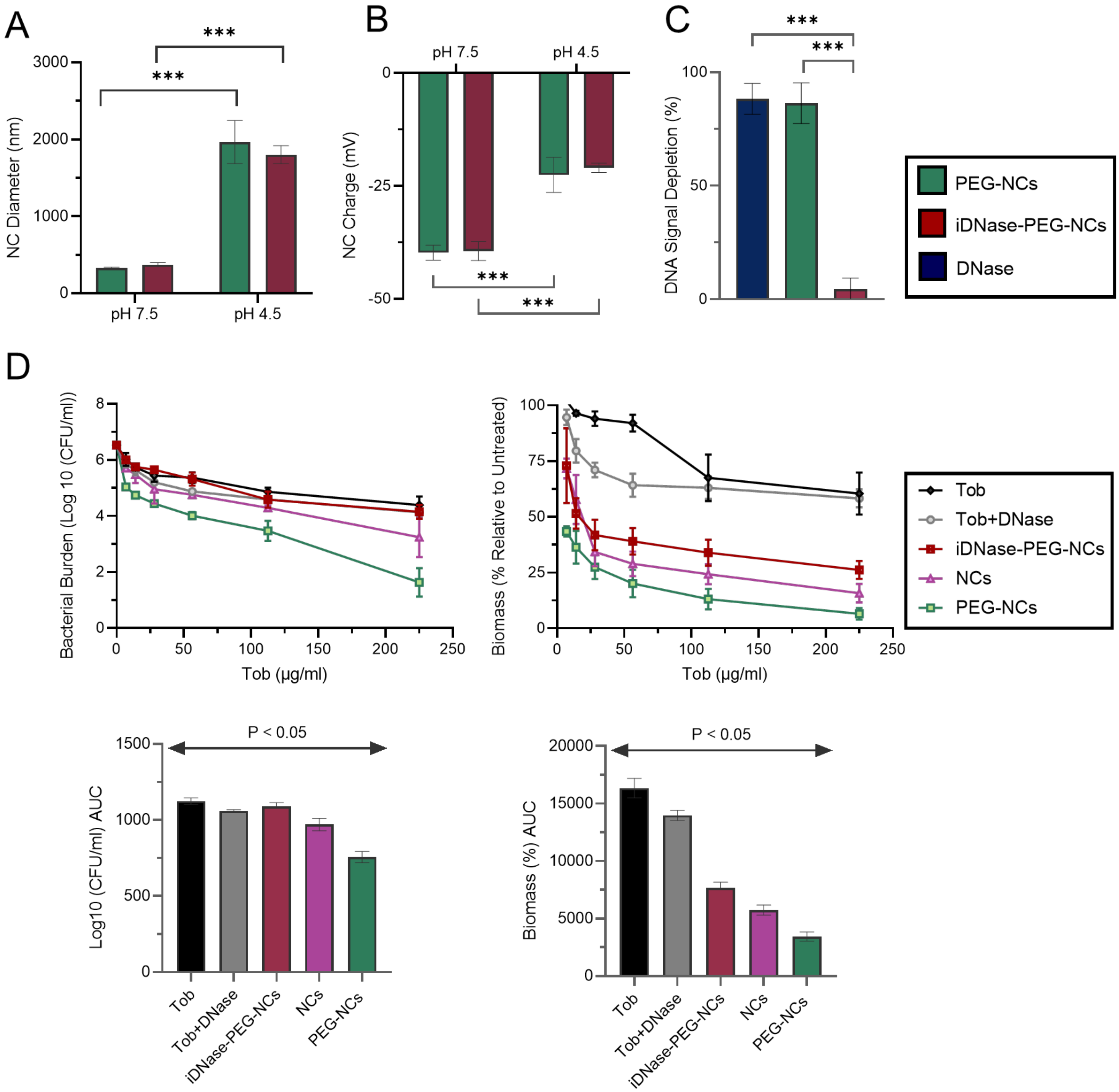


**Figure S5. DNase inactivation preserves nanocomplex physicochemical properties but reduces antibiofilm activity.** Inactive DNase PEG-NCs (iDNase-PEG-NCs) were prepared as a control to assess the contribution of DNase activity. (A) Hydrodynamic diameter, (B) zeta potential, and (C) DNase activity of PEG-NCs and iDNase-PEG-NCs. (D) Concentration-dependent bacterial burden and biofilm biomass in established LESB58 biofilms after treatment with Tob+DNase, iDNase-PEG-NCs, NCs, or PEG-NCs, with corresponding area-under-the-curve (AUC) analyses. Data for Tob+DNase, NCs, and PEG-NCs are also presented in Figure 3A,B of the main text, with iDNase-PEG-NCs included here for comparison. AUC values were compared using one-way ANOVA followed by Tukey's multiple-comparisons test. Double-headed arrows indicate statistically significant pairwise differences between the connected treatment groups based on Tukey-adjusted p-values ($P < 0.05$). Data are mean ± SD from three independent experiments. ***$P < 0.001$.

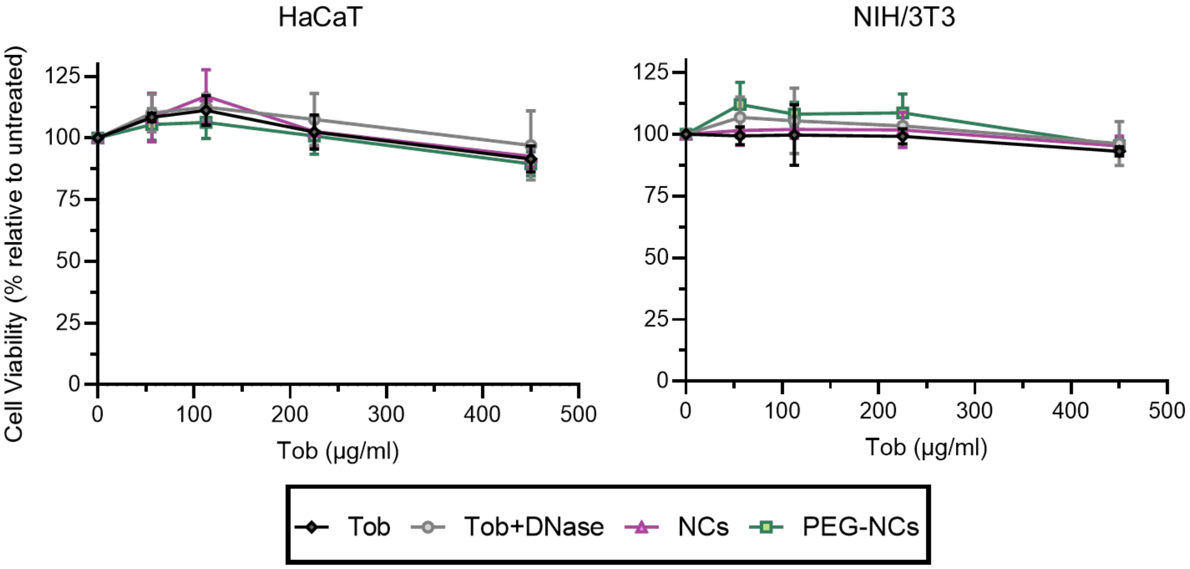


**Figure S6. Tob, Tob+DNase, NCs, and PEG-NCs are biocompatible with HaCaT and NIH/3T3 cells in vitro.** 10,000 cells were exposed to increasing Tob-equivalent concentrations of Tob, Tob+DNase, NCs, or PEG-NCs for 24 h. HaCaT and NIH/3T3 cell viability measured using the resazurin assay and expressed relative to untreated cells. Data are presented as mean ± SD from three independent experiments.

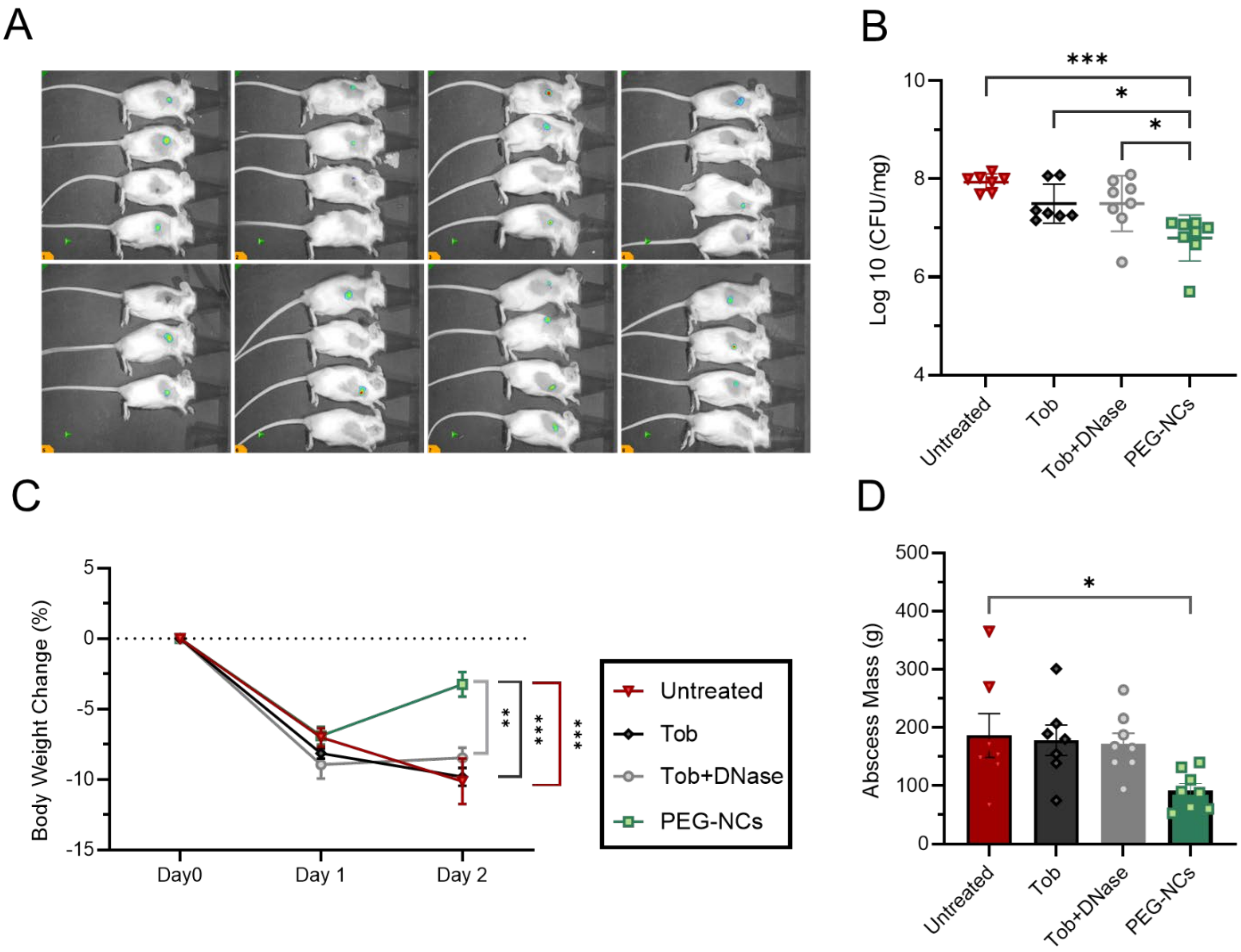


**Figure S7. PEG-NCs reduce bacterial burden in the independent 24-h murine abscess cohort.** Mice were infected on Day 0, treated on Day 1, and sacrificed on Day 2. (A) Representative IVIS images acquired 24 h after treatment. IVIS imaging was used to assess infection localization and not as a direct measure of bacterial burden. (B) Viable bacterial burden expressed as log 10 CFU/mg. (C) Body-weight change relative to Day 0. (D) Abscess mass at collection. Each mouse represents one biological replicate (n=8 mice per group). Individual mice and mean ± SD are shown in B and D; longitudinal data are presented as mean ± SEM. *P<0.05 and ***P<0.001.

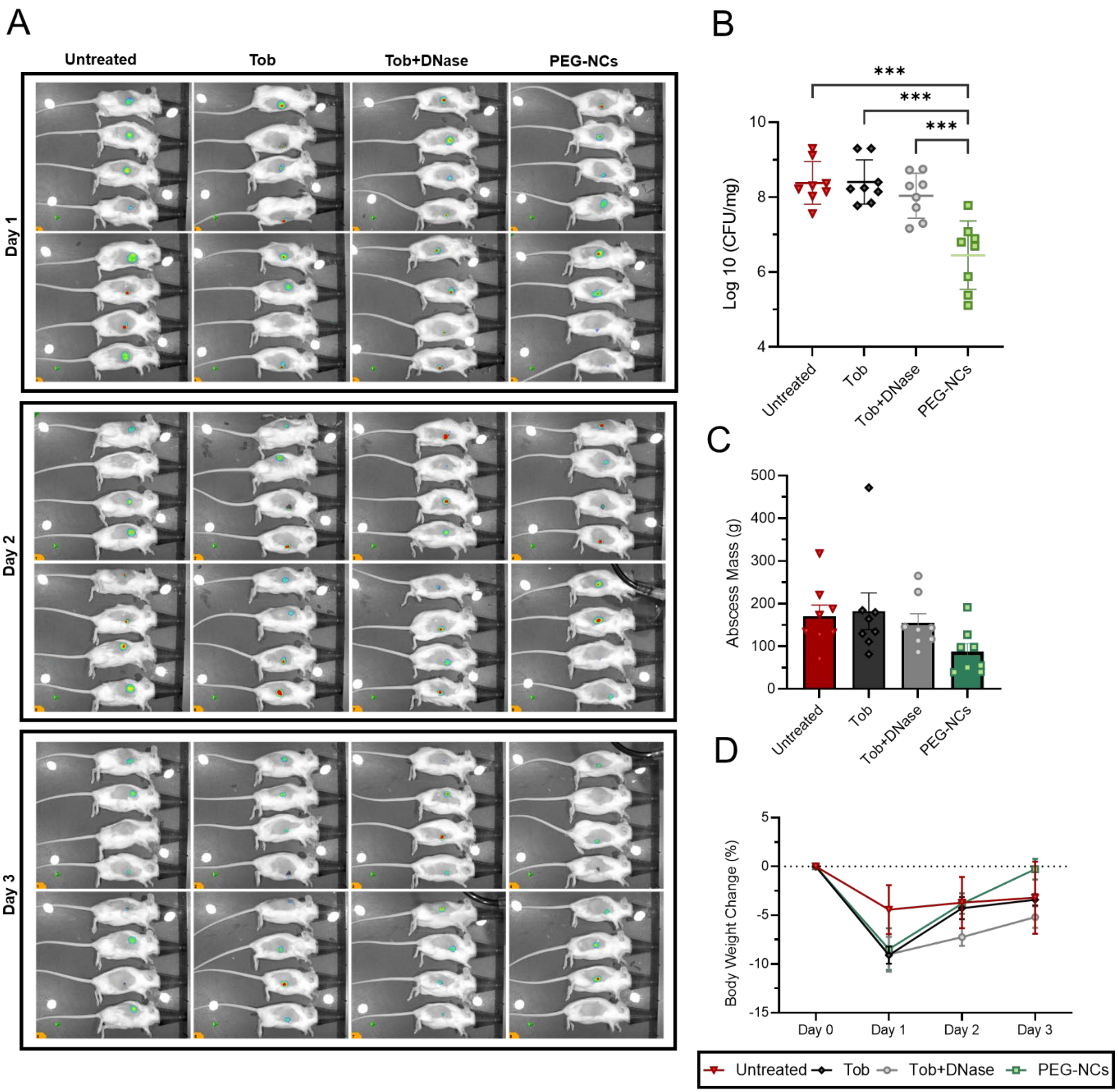


**Figure S8. PEG-NCs reduce bacterial burden in the independent 48-h murine abscess cohort.** Mice were infected on Day 0, treated on Day 1, and sacrificed on Day 3. (A) Representative IVIS images acquired before treatment and at 24 and 48 h after treatment. IVIS imaging was used to track the infection localization, not as a direct measure of bacterial burden. (B) Viable bacterial burden expressed as log10 CFU/mg. (C) Abscess mass at collection. (D) Body-weight change relative to Day 0. Each mouse represents one biological replicate ((n=8) mice per group). Individual mice and mean ± SD are shown in B and C; longitudinal data are presented as mean ± SEM. ***P<0.001.

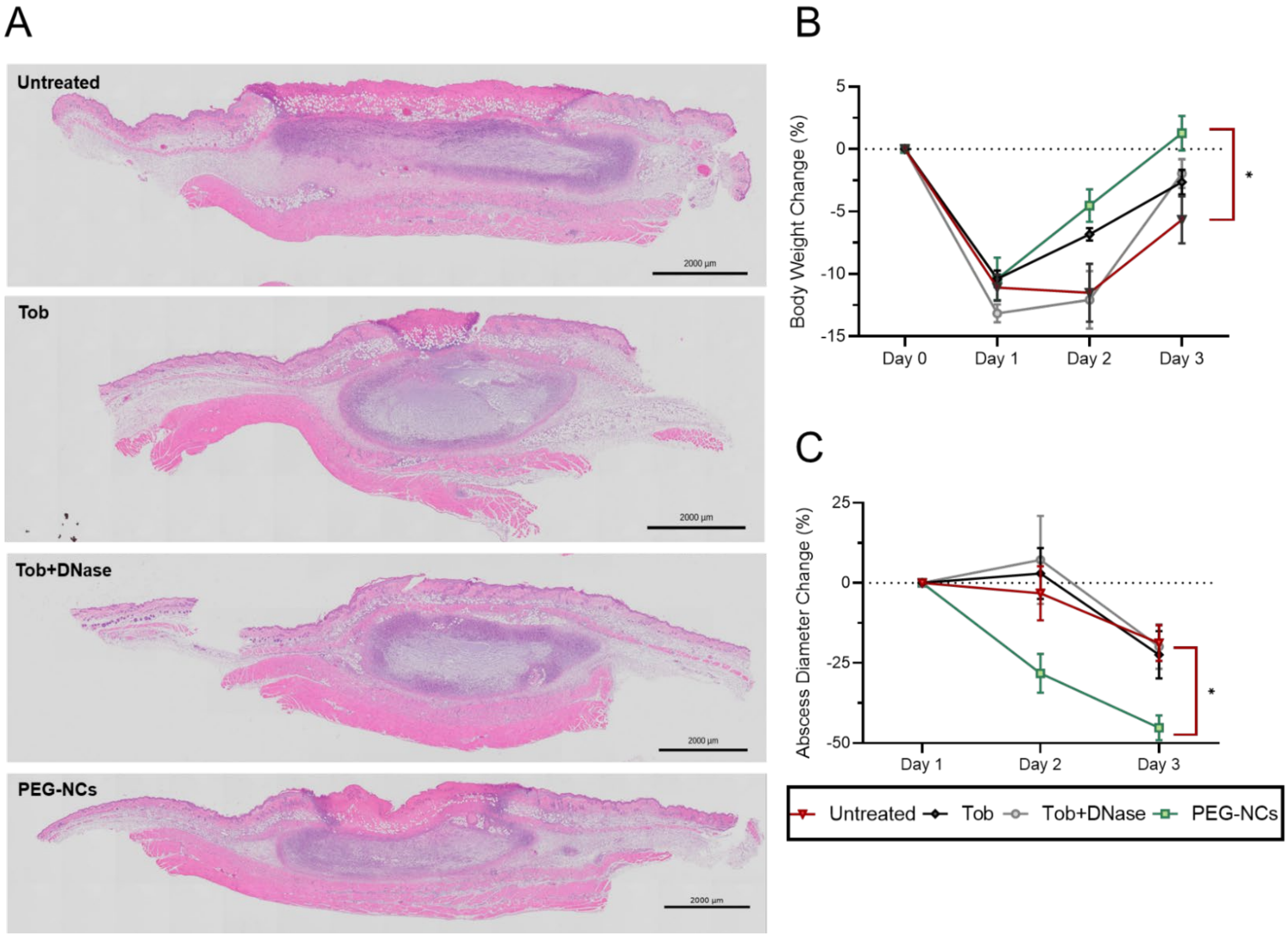


**Figure S9. Longitudinal outcomes and representative histology from the independent 48-h murine abscess cohort.** Mice were infected on Day 0, treated on Day 1, and sacrificed on Day 3. (A) Representative H&E-stained abscess sections, scale bar=2000 µm. (B) Body weight change relative to Day 0. (C) Abscess-diameter change relative to the pretreatment measurement on Day 1. Data are presented as mean ± SEM (n=4 mice per group). * $P < 0.05$ between untreated and PEG-NC treatment groups.
